# Non-canonical cytokinesis driven by mechanical uncoupling via nematic flows and adhesion-based invagination

**DOI:** 10.1101/2025.10.15.682552

**Authors:** Xin Tong, Yuting I. Li, Joséphine Schelle, Edouard Hannezo, Carl-Philipp Heisenberg

## Abstract

Cleavage - the series of rapid cell divisions that follow fertilization - marks the onset of metazoan development and represents a deeply conserved evolutionary process. Across animals, two principal modes exist: complete (holoblastic) and incomplete (meroblastic) cleavage. While holoblastic cleavage resembles conventional cytokinesis both *in vitro* and *in vivo*, the mechanisms underlying meroblastic cleavage have remained poorly understood. Using zebrafish embryos as a model, we show that meroblastic cleavage proceeds through a distinct two-step mechanism. The process begins with the assembly and contraction of a large, arc-shaped actomyosin cable. However, this contractile event alone is insufficient to complete division. A second phase, driven by cadherin-mediated membrane adhesion, is required to invaginate the furrow ridge. Strikingly, this transition depends on mechanical uncoupling of the contractile cable from the surrounding cortex. We demonstrate that such uncoupling arises from an active nematic instability, which both enhances contractility along the cable and generates actin depletion zones that relieve lateral connections. Together, these findings reveal that meroblastic cleavage is governed not by a single actomyosin-based event but by a sequential interplay between cytoskeletal contraction and cadherin-dependent adhesion, highlighting a mechanism fundamentally distinct from canonical cytokinesis.

---

Cytokinesis is the final step of cell division, during which a cleavage furrow partitions the cytoplasm and segregates the duplicated chromosomes into distinct daughter cells. This process has been extensively studied in cells where cytokinesis proceeds to completion^1–3^. There, cytokinesis is initiated by the assembly of a contractile ring at the future site of the cleavage furrow. This ring is composed of actin filaments, myosin-2 motors, and regulatory proteins such as formin, anillin, and septin^3–8^. Contraction of the actomyosin ring - resembling a ‘purse-string’ mechanism - drives ring constriction, pulling the plasma membrane inward to form the cleavage furrow and ultimately leading to the physical separation of the daughter cells^9–12^. While this widely accepted model effectively explains complete cytokinesis both *in vitro* and *in vivo*, the mechanistic basis of alternative division modes, such as those involving incomplete cytokinesis, remains poorly understood.

In most animals, embryonic development begins with the fertilized egg (zygote) undergoing several rounds of reductive cell divisions, known as cleavages, which partition the large volume of the egg into smaller cells that form the first embryonic tissue, the blastoderm^13–15^. These embryonic cleavages can be either complete (holoblastic), fully separating the daughter cells, or incomplete (meroblastic), only partially dividing them. Meroblastic cleavages typically occur in organisms whose eggs contain a large proportion of dense yolk granules, such as fish, birds and reptiles, while holoblastic cleavages are found in organisms with sparse or no yolk in their eggs, such as echinoderms, amphibians and most mammals^13,14^.

Zebrafish embryos serve as a classic model for studying meroblastic cleavage^16–23^ (Movie S1). In zebrafish, the initial cleavage divisions are associated with the formation of an arc-shaped actomyosin cable at the animal pole of the zygote^24,25^. This structure has been proposed to facilitate cleavage furrow formation and the partial separation of the zygote’s animal region - the blastodisc - which contains most of the cytoplasm destined to give rise to the embryo proper^14,26,27^. However, the mechanistic basis of cleavage furrow formation and indentation, as well as the specific role of the actomyosin cable in this process, remains unresolved.

From a mechanical perspective, meroblastic cleavage in large eggs presents distinct challenges. First, in both meroblastic and holoblastic contexts, successful cytokinesis requires that the inward forces generated by the contractile actin cable or ring exceed the opposing cortical tensions exerted from the cell poles^9,11,28–31^. However, as cell size increases, the opposing cortical tension grows faster than the inward contractile force that drives furrow ingression^9^. Consequently, larger eggs demand substantially higher contractility than smaller cells to overcome this geometric constraint. Second, in holoblastic cleavage, complete actomyosin rings progressively constrict, increasing curvature and thereby amplifying inward forces - a positive feedback mechanism that promotes ring closure^9,32,33^. In meroblastic cleavage, however, this effect is diminished or even inverted: the contractile cable flattens over time, reducing curvature and hindering closure. Collectively, these different mechanical considerations suggest that the strategies enabling meroblastic cleavage must differ fundamentally from those driving complete cell division.

## The first meroblastic cleavages can be divided into two consecutive phases

To investigate how meroblastic cleavages in zebrafish are mechanistically achieved, we examined the dynamic morphological changes that the embryo (zygote) undergoes during its first cleavage. Performing quantitative reconstructions and morphological shape analysis on Tg(*actb2:mCherry-CAAX*) embryos with fluorescently labelled plasma membranes, we identified two distinct phases of cytokinesis (Fig. 1A, 1A’, 1B, 1C, S1B, Movie S2, S3, S4). In the first phase (Phase 1), the cleavage furrow began to form at the center of the animal pole and extended circumferentially toward the vegetal pole (Fig. 1B, 1D). This furrow extension progressed until it reached the yolky region of the zygote. During this time, the furrow gradually deepened, accompanied by only a modest reduction in curvature (Fig. 1E, S1D, S1F). Phase 1 was followed by a second phase (Phase 2), which commenced at the onset of mitosis of the second cell cycle (prior to the second cleavage), when furrow extension had ceased (Fig. 1B). During this second phase, the plasma membrane began to invaginate along the ridge of the existing furrow, forming an adhesive membrane fold (septum) that further separated the two daughter blastomeres. The septum continued to invaginate until it reached the yolky region by the end of mitosis (Fig. 1B, 1D). Notably, during Phase 2, the depth, length and curvature of the furrow ridge remained largely unchanged (Fig. 1E, 1F, 1F’, S1C, S1D, S1E). This suggests that Phases 1 and 2 correspond to two distinct morphogenetic processes: furrow indentation and furrow invagination, respectively (Fig. 1C).

**Figure 1.**
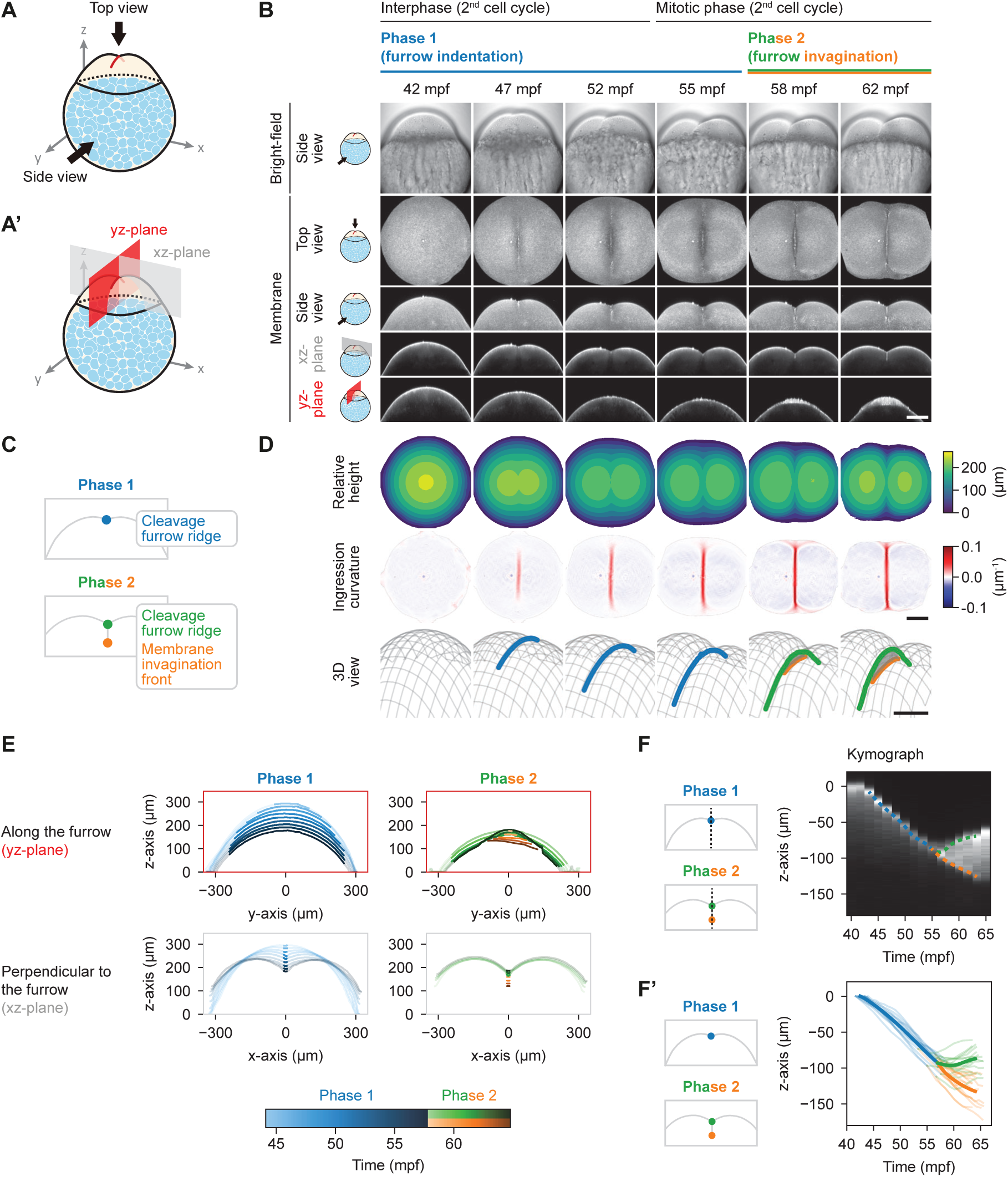
Dynamic changes in cleavage furrow geometry during the 1st cleavage division. (A and A’) Schematics of imaging orientations, illustrating the top view (toward the animal pole, A), the side view (along the y-axis, A) and two sectional views (the yz-plane sectioning along the cleavage furrow and the xz-plane sectioning perpendicular to the cleavage furrow, A’). (B) Bright-field (upper row) and fluorescence images (middle and lower row) of a Tg(*actb2:mCherry-CAAX*) zebrafish embryo with labelled plasma membrane at consecutive time points during the 1st cleavage of the fertilized egg (zygote). The different orientations are as indicated on the left and in (A, A’). The top and side views are maximum intensity projections. Scale bar, 150 μm. (C) Schematics of the two consecutive phases (Phase 1 and Phase 2) during the 1st cleavage of the zygote. Blue illustrates the furrow ridge during Phase 1, green the furrow ridge (contact point) during Phase 2 and orange the leading edge of the invaginating membrane septum during Phase 2. (D) Height maps (upper row), ingression curvature (*k*_1_ principal curvature; middle row) and 3D renderings (lower row) of the animal surface of the zygote at consecutive time points during the 1st cleavage as shown in (B). Scale bar, 150 μm. (E) Sectional line views of the animal surface of the zygote at consecutive time points during Phase 1 and Phase 2, color-coded as shown in the the color bar below - note that during Phase 2, the green lines indicate the surface indentation, while the orange lines mark the leading edge of the invaginating septum. The upper row panels are along the furrow and the lower row panel perpendicular to it. The transparent part of the lines indicates the outline of the zygote on the sectional plane and the bold part indicates the region with positive ingression curvature (the position of the geometrical furrow ridge). (F) Kymograph of the furrow indentation and membrane invagination along the z-axis at the embryo center (black dotted line in the left schematics), during the 1st cleavage division of a Tg(*actb2:mCherry-CAAX*) embryo. The blue, green and orange dotted lines correspond to features in (C). (F’) Line plot showing the positions of the cleavage furrow ridge and the membrane invagination front as a function of time during the 1st cleavage of the zygote. Color illustrations as in (C). N = 15 clutches from 8 different days, n = 15 embryos.

## Convergent flows of actin filaments drive actin cable formation at the cleavage furrow

To investigate the mechanochemical processes underlying the first meroblastic cleavage, we focused on the cortical actomyosin network, which has previously been implicated in cytokinesis of animal cells^34,35^. Specifically, we examined how this network reorganizes when the cleavage furrow initiates and begins to extend. Using Tg(*actb2:Utrophin-mCherry*) embryos with fluorescently labelled actin, we observed that furrow formation was marked by a striking reorganization of the actin cytoskeleton (Fig. 2A, 2B, Movie S5). Initially, actin was distributed as an isotropic, punctate network at the animal pole (Fig. 2C, Movie S7). As cleavage progressed, this network gradually transformed into a highly aligned, cable-like structure along the forming furrow (Fig. 2C).

**Figure 2.**
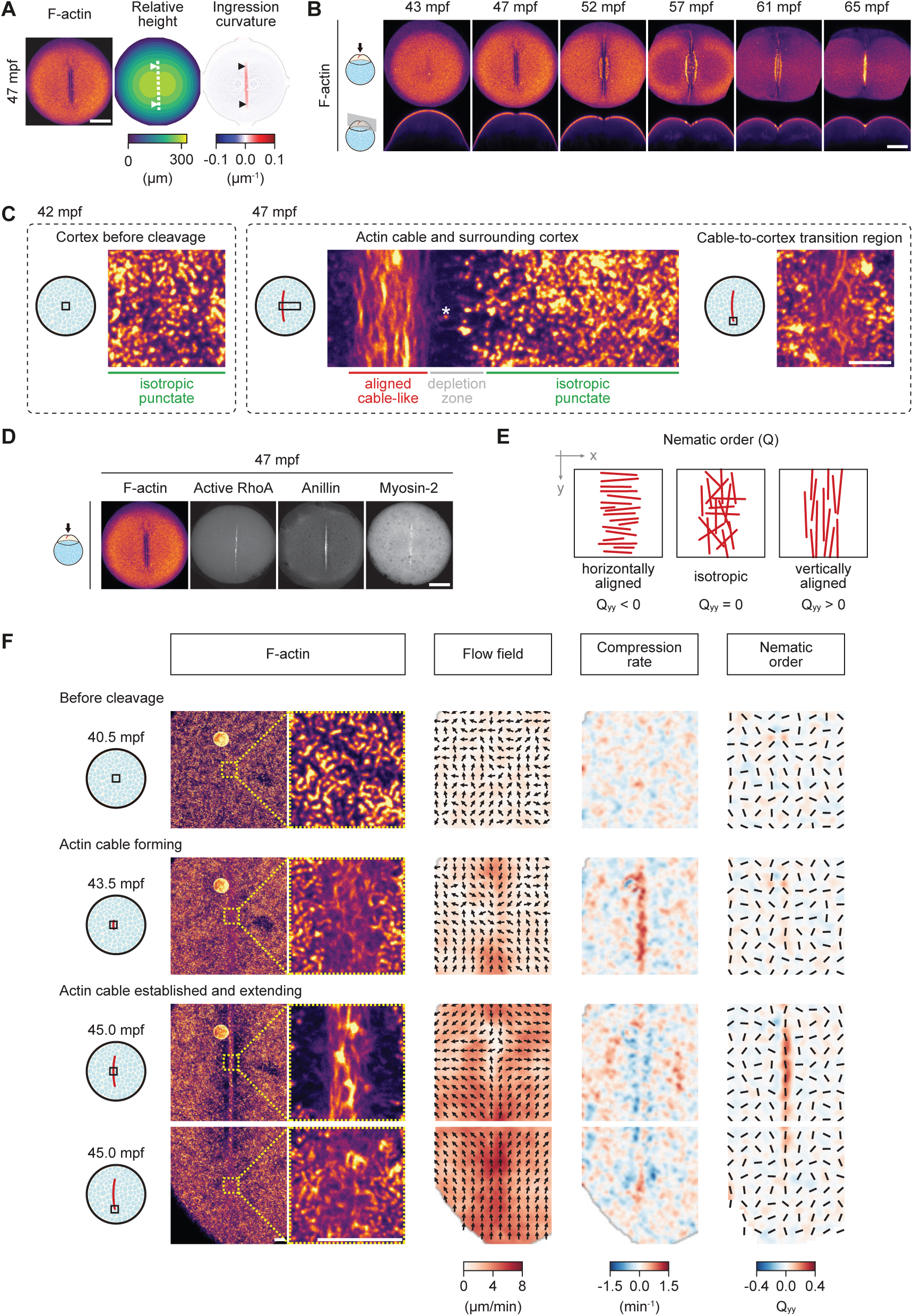
Dynamic changes of the actin cortex during the 1st cleavage. (A) Fluorescence images of a Tg(*actb2:Utrophin-mCherry*) embryo with labelled F-actin (left), its surface height map (middle) and surface ingression curvature (*k*_1_ principal curvature; right) at 47 mpf. Arrow heads indicate the two termini of the actin cable, and dotted lines the position of the furrow ridge determined by positive surface ingression curvature. Scale bar, 150 μm. (B) Fluorescence images of a Tg(*actb2:Utrophin-mCherry*) embryo with labelled F-actin at consecutive time points during the 1st cleavage division. The different views are as indicated on the left and in Fig. 1A, 1A’. Top views are maximum intensity projections. Scale bar, 150 μm. (C) High-resolution fluorescence images of a Tg(*actb2:Lifeact-EGFP*) embryo with maximum intensity projection of a 15 μm z-stack showing the organizations of F-actin before (left panel) and during (right panel) the 1st cleavage division. The imaging positions are indicated by the accompanying schematics. Scale bar, 5 μm. (D) Fluorescence images of a Tg(*actb2:Utrophin-mcherry*) embryo with labelled F-actin, a wild-type embryo injected with 1 μg mRNA of mNeonGreen-2xrGBD labelling active RhoA, a Tg(*actb2:anillin-mNeonGreen*) embryo with labelled Anillin, and a Tg(*actb2:Myl12.1-EGFP*) with labelled myosin-2 light chain at 47 mpf. Top views are maximum intensity projections. Scale bar, 150 μm. (E) Schematics of nematic order (Q-tensor). Positive Q_yy_ value indicates alignments along the y-axis and negative Q_yy_ value along the x-axis. (F) Fluorescence images of a Tg(*actb2:Lifeact-EGFP*) embryo with labelled cortical actin and their zoom-in views (1st column) at three representative time points during the formation of the actin cable, with the corresponding flow field (2nd column; arrows indicating the local flow direction and color-coding the local velocity magnitude), compression rate (3rd column; red indicating compression and blue expansion) and nematic order (4th column; bars indicate the orientation of alignment and color-coding the value of Q_yy_ component; red indicates the alignment along y-axis and blue along x-axis). The position and time points are shown by the schematics on the left. Scale bar, 10 μm.

The onset of actin cable formation coincided with a localized upregulation of the small Rho GTPase RhoA, marking the region where the cable was assembled, as observed in embryos injected with mRNA encoding the active RhoA sensor mNeonGreen-2xrGBD^36^ (Fig. 2D, S2B). This suggests that, as in complete cytokinesis in other systems, signals emanating from the growing mitotic spindle midzone locally activate RhoA at the center of the animal pole^37–40^. In turn, RhoA activates formin and non-muscle myosin-2, facilitating the recruitment and alignment of actin filaments into a contractile cable that drives cleavage furrow initiation and cytokinesis^34,41–43^. Consistent with this mechanism, we observed a strong accumulation of myosin-2 within the forming actin cable, along with the recruitment of additional actin-binding proteins typically associated with the cleavage furrow in other cell types, including anillin^7,44,45^, as visualized in Tg(*actb2:myl12.1-EGFP*) and Tg(*actb2:anillin-mNeonGreen*) embryos (Fig. 2D, S2C, S2D, S3A). Together, these findings indicate that actin cable formation during meroblastic cleavage engages a set of conserved biochemical effectors, akin to those operating in complete cytokinesis.

Surprisingly, while the actin cable remained connected with the actin cortex at its ends, it became laterally detached, as evidenced by the emergence of regions devoid of discernible actin, which we refer to as *depletion zones* (Fig. 2C). This organization contrasts sharply with classical systems, where the actomyosin ring remains continuously connected to the actin cortex along both its ends and sides^46–48^. High spatiotemporal-resolution imaging revealed that this reorganization was driven by convergent actin flows (Fig. 2F, 2nd row, Movie S8). These flows first emerged at the center of the animal pole, coinciding with the site of furrow initiation, and progressively extended toward both ends of the elongating furrow (Fig. 2F, 2nd row). Notably, these flows were much shorter-ranged compared to classical cytokinesis, such as in *C. elegans* early cleavages, where these flows extend to the poles of the dividing cell^32,47,49^. Furthermore, unlike classical cytokinesis, once a well-defined actin cable with high nematic order had formed, the short-range convergent flows ceased, and were replaced by long-range divergent flows (Fig. 2F, 3rd row). Collectively, these findings suggest that the actin rearrangements underlying cleavage furrow formation in this system differ fundamentally from those observed in classical models of cytokinesis.

To interpret the observed dynamics, we constructed a physical model grounded in active gel theory^50,51^. Previous models of cytokinesis generally predicted inward actin flows both at the ends of the forming actin cable and along its lateral sides^9,52^. These predictions, however, are inconsistent with our experimental findings, which reveal outward divergent flows along the cable’s sides and inward flows at its ends (Fig. 2F, 3rd row). Furthermore, theoretical frameworks that are limited to isotropic contractility did not predict the experimentally observed actin depletion zones within a physiologically relevant parameter regime (see SI Theory Note for details). We therefore sought to identify the minimal additional features required for the theory to accurately reproduce our experimental observations, progressively increasing model complexity as needed.

As detailed in the SI Theory Note, this approach led us to incorporate two key physical properties into the model. First, nematic contractility, which produces stronger contraction along the axis of aligned actin filaments (Fig. 3B). Second, nematic order stabilization, a hallmark of anisotropic structures, which maintains the cable-like architecture once formed^53–57^ (see SI Theory Note for details; Fig. S3B, S3B’, Movie S9). Guided by our experimental data, we modelled the spindle midzone as a region of myosin-2 activation that drives the formation of a contractile actomyosin network, while the surrounding cortex was represented as punctate actin with minimal myosin-2 activity (Fig. 3A, S3A, and SI Theory Note).

**Figure 3.**
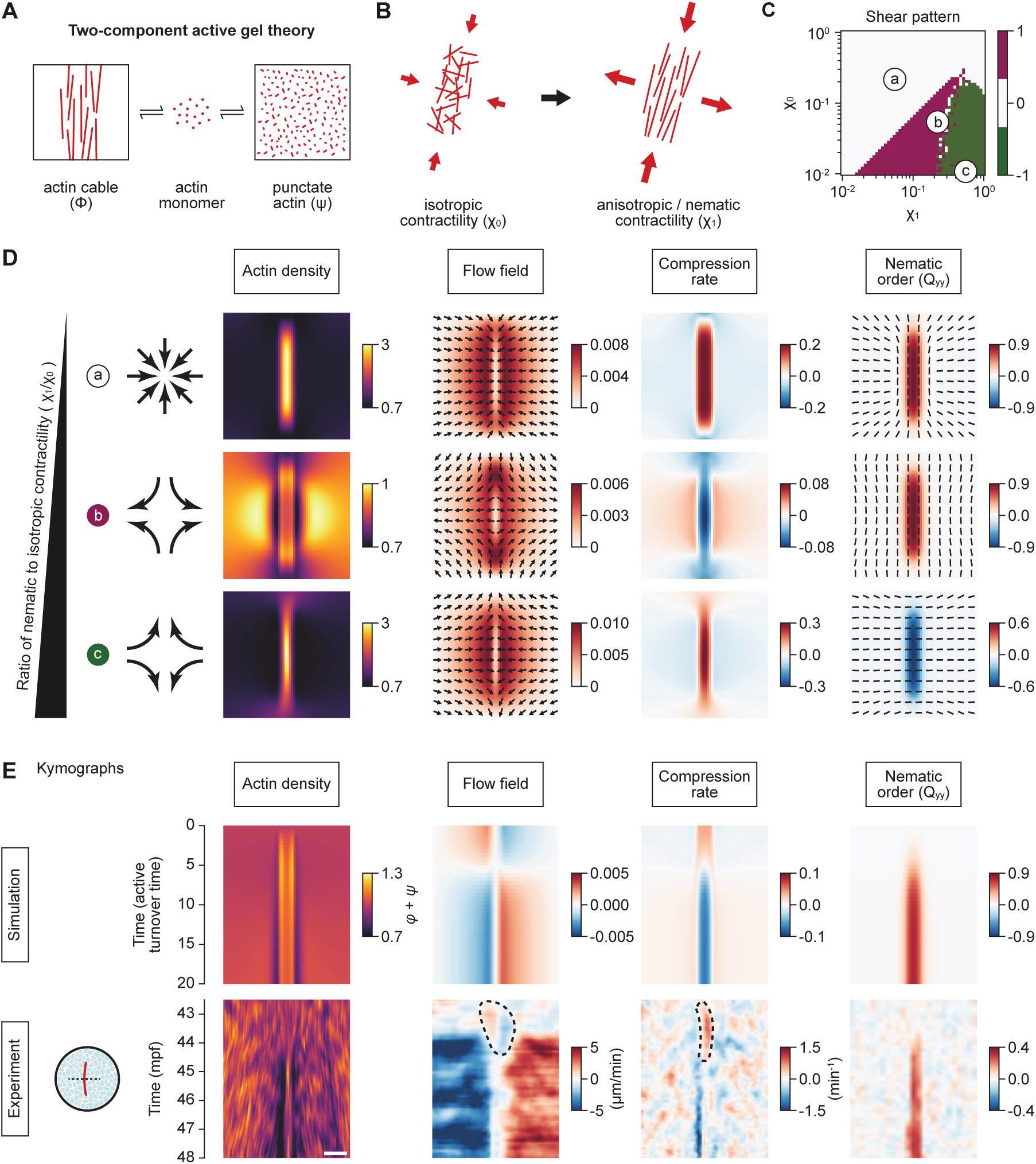
Two-component active gel theory capturing dynamic changes of the actin cortex during the 1st cleavage. (A) Schematics illustrating the two-component active gel theory. (B) Schematics illustrating isotropic contraction (*χ*_0_) and anisotropic contraction (*χ*_1_). (C) Phase diagram of the shear flow patterns as a function of isotropic contractility (*χ*_0_) and anisotropic contractility (*χ*_1_). (D) Simulations of the three scenarios of shear patterns in (C) as *χ*_1_/*χ*_0_ increases, shown by their corresponding actin density field (1st column), flow field (2nd column; arrows indicate the local flow direction and color-coding the local velocity magnitude), compression rate (3rd column; red indicates compression and blue divergence) and nematic order (4th column; bars indicate the orientation of alignment and color-coding the value of Q_yy_ component; red indicates the alignment along y-axis and blue along x-axis). (E) Kymographs of actin density field (1st column), flow field (2nd column; red indicates moving to the right and blue to the left), compression rate (3rd column; red indicates compression and blue divergence) and nematic order (4th column; red indicates the alignment along y-axis and blue along x-axis) along the middle line perpendicular to the actin cable in simulations (upper row) and experimental data (lower row). Scale bar, 20 μm.

Our model uncovered distinct classes of flow fields that depend on the ratio of isotropic to nematic contractility (Fig. 3C). When nematic contractility was low, the model predicted globally isotropic flows toward the forming actin cable without generating a depletion zone, consistent with classical cytokinesis^47,52,58,59^ (Fig. 3D, region a). By contrast, when nematic contractility exceeded isotropic contractility, the predicted dynamics closely matched our experimental observations (Fig. 3D, region b). In this regime, convergent flows initially concentrated and aligned actin filaments into a central cable (Fig. 3E, upper panel). Nematic contractility within the cable then drove inward flows at its ends and converted the initial convergent flows at its lateral sides into outward divergent flows (Fig. 3E, upper panel). These dynamics not only explained cable formation but also predicted the emergence of lateral actin depletion zones, effectively decoupling the cable from the surrounding cortex (Fig. 3E, lower panel). At very high nematic contractilities, the model predicted cable instability (Fig. 3D, region c), with nematic order aligning perpendicular to the cable and producing lateral inflows coupled to outflows at the ends. Together, these findings suggest that the ratio of nematic to isotropic contractility - specifically the regime represented by region b in our phase diagram (Fig. 3C) - regulated by signals from the growing spindle midzone are sufficient to account for the cortical actin rearrangements and flow patterns observed during the first cleavage division.

To experimentally challenge these predictions, we first examined whether the actin cable is contractile and whether its tension scales with the degree of filament alignment. We performed UV-laser cuts of the actin cable either perpendicular or parallel to its axis and used initial recoil velocities as a proxy for tension^60,61^. Recoil was predominantly observed following perpendicular cuts (Fig. 4A, 4A’, Movie S10), indicating that the cable primarily contracts along its length, as predicted by region b of our phase diagram (Fig. 3C). Moreover, this anisotropy in tension increased with the degree of nematic order during cable formation (Fig. 4B, 4B’, S4B, S4B’), consistent with the model assumption that filament alignment enhances both the strength and directionality of contractile tension. As a control, we also performed laser ablations of the cortex both parallel and perpendicular to the actin cable (Fig. 4A, 4A’). In both orientations, we detected minimal recoil upon cutting, in line with our observation that myosin-2 is localized predominantly in the cable region (Fig. S3A) and our model assumption of two actomyosin populations.

**Figure 4.**
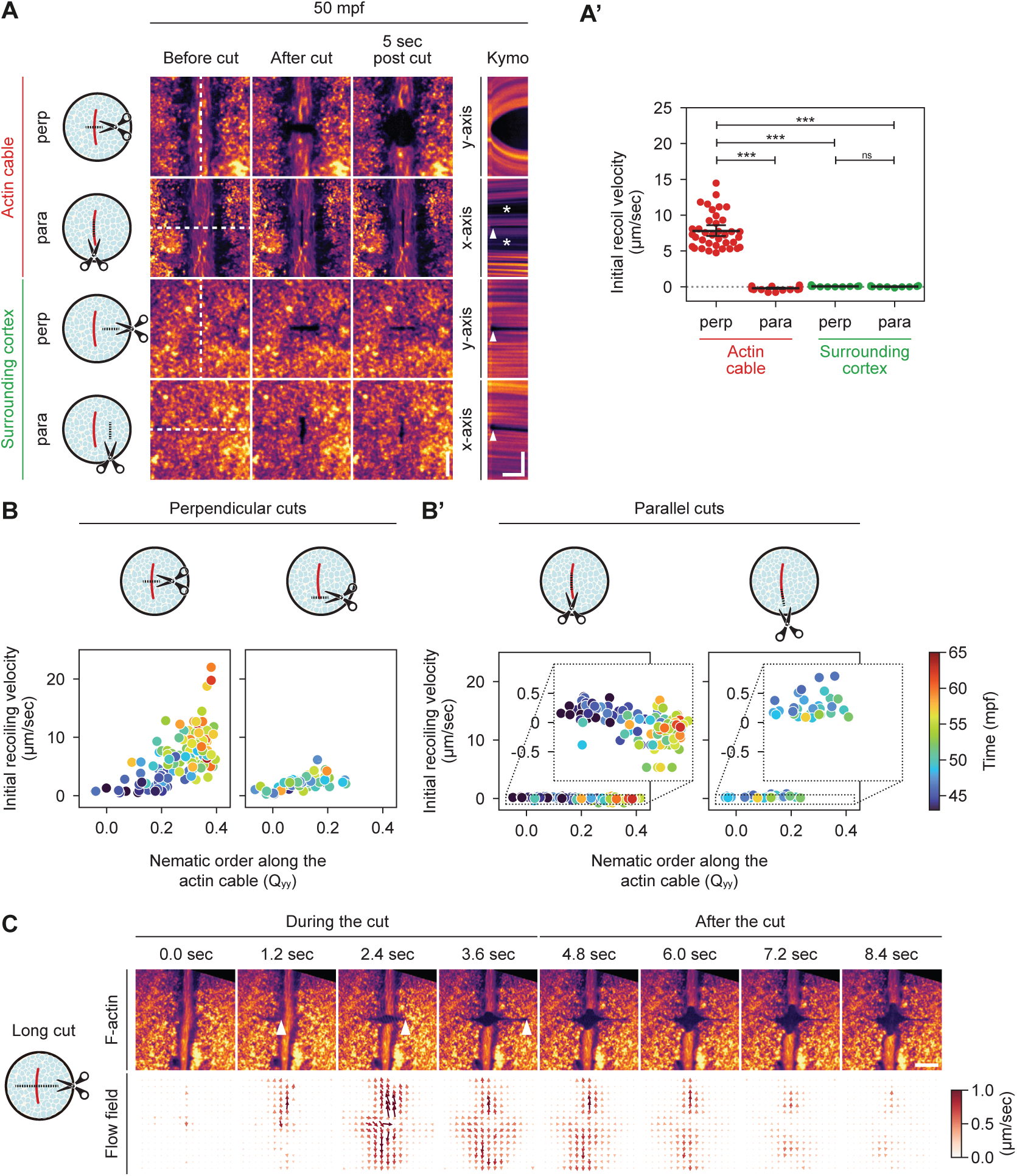
Tension measurements of the actomyosin cable and adjacent cortex during the 1st cleavage. (A) Fluorescence images of UV-laser ablations of the actin cable or surrounding cortex in Tg(*actb2:Lifeact-EGFP*) embryos with labelled F-actin at 50 mpf. The cut positions and orientations are shown by the schematics on the left. The fluorescence images before cut, 0 and 5 seconds after cut are shown on the left three columns, respectively. Kymographs along y-axis and x-axis, as outlined by the white dashed lines, are shown in the right column. The arrow heads indicate the position of the cut and asterisks the actin deletion zones. Horizontal scale bar, 5 sec and vertical scale bar, 10 μm. (A’) Plot of initial recoil velocities of the actin cable and surrounding cortex after UV-laser ablations for the cut positions and orientations as shown in (A). Data are shown in a categorical scatter plot with the bold horizontal bars representing the means and the 95% confidence intervals. Perpendicular, actin cable: N = 13 clutches from 5 different days, n = 37 cuts on 31 embryos; parallel, actin cable: N = 8 clutches from 4 different days, n = 28 cuts on 28 embryos; perpendicular, surrounding cortex: N = 5 clutches from 2 different days, n = 11 cuts on 11 embryos; parallel, surrounding cortex: N = 5 clutches from 2 different days, n = 12 cuts on 12 embryos. Statistical tests, Kruskal-Wallis test followed by post-hoc Wilcoxon rank-sum tests with Bonferroni correction: cable perp and cable para, \*\*\**p* < 0.001; cable perp and cortex perp, \*\*\**p* < 0.001; cable perp and cortex para, \*\*\**p* < 0.001; cortex perp and cortex para, *p* = 1. (B, B’) Scatter plots of the nematic order on the actin cable versus the initial recoil velocities after UV-laser ablations during the 1st cleavage division. The ablations are done in the positions and orientations as shown by the corresponding schematics above. The time of the ablations are color-coded. Perpendicular, mature actin cable: N = 14 clutches from 5 different days, n = 134 cuts on 87 embryos; perpendicular, actin cable termini: N = 12 clutches from 5 different days, n = 49 cuts on 39 embryos; parallel, mature actin cable: N = 10 clutches from 5 different days, n = 102 cuts on 59 embryos; parallel, actin cable termini: N = 8 clutches from 4 different days, n = 34 cuts on 33 embryos. (C) Fluorescence images of an exemplary 80 μm-long UV-laser cut perpendicular to the actin cable in a Tg(*actb2:Lifeact-EGFP*) embryo at consecutive time points during and after ablation (upper row), and their corresponding flow fields (lower row) at 50 mpf. The arrow heads on the upper row indicate the trajectory of the UV-laser during cutting. The magnitude of local velocities on the bottom row are indicated by both the arrow length and color-coding (as shown in the right color bar). Scale bar, 20 μm. N = 5 clutches from 4 different days, n = 6 embryos.

A key prediction of our model is the emergence of actin depletion zones at the lateral edges of the cable, which are expected to mechanically uncouple the cable from the cortex. To directly test this, we performed large-scale laser ablations of both the cable and the adjacent cortex and monitored the extent to which recoil of the cable is accompanied by corresponding movements in the cortex, serving as a readout for mechanical coupling between the two actin networks (Fig. 4C, Movie S12). While pronounced recoil of the cable upon laser ablation was observed, only minimal recoil was observed in the adjacent cortex, supporting the notion that the actin depletion zones at the lateral sides of the cable mechanically uncouple it from the surrounding cortex.

To further investigate the role of convergent actin flows in cable assembly and tension generation, we disrupted these flows by overexpressing a constitutively active (CA) form of RhoA - a key regulator of actin network organization and myosin motor activity^6,38,40–43^. Zygotes injected with CARhoA mRNA exhibited a spectrum of phenotypes of varying severity, likely reflecting differences in CARhoA expression levels (Fig. 5A, 5D, Movie S13, S14). In general, CARhoA overexpression altered actin organization, resulting in less pronounced filament alignment within the cable and reduced formation of the actin depletion zone (Fig. S5B, S5D). Moreover, the isotropic contractility of the cortex was slightly increased, as revealed by laser ablation experiments (Fig. 5B, S5F). Despite these alterations, the relationship between recoil velocity and local nematic order within the actin cable remained similar to that observed under wild-type conditions (Fig. 5C). This indicates that CARhoA overexpression does not affect nematic contractility per se, but instead lowers the maximum attainable level of nematic order (Fig. 5E, S5B, S5E).

**Figure 5.**
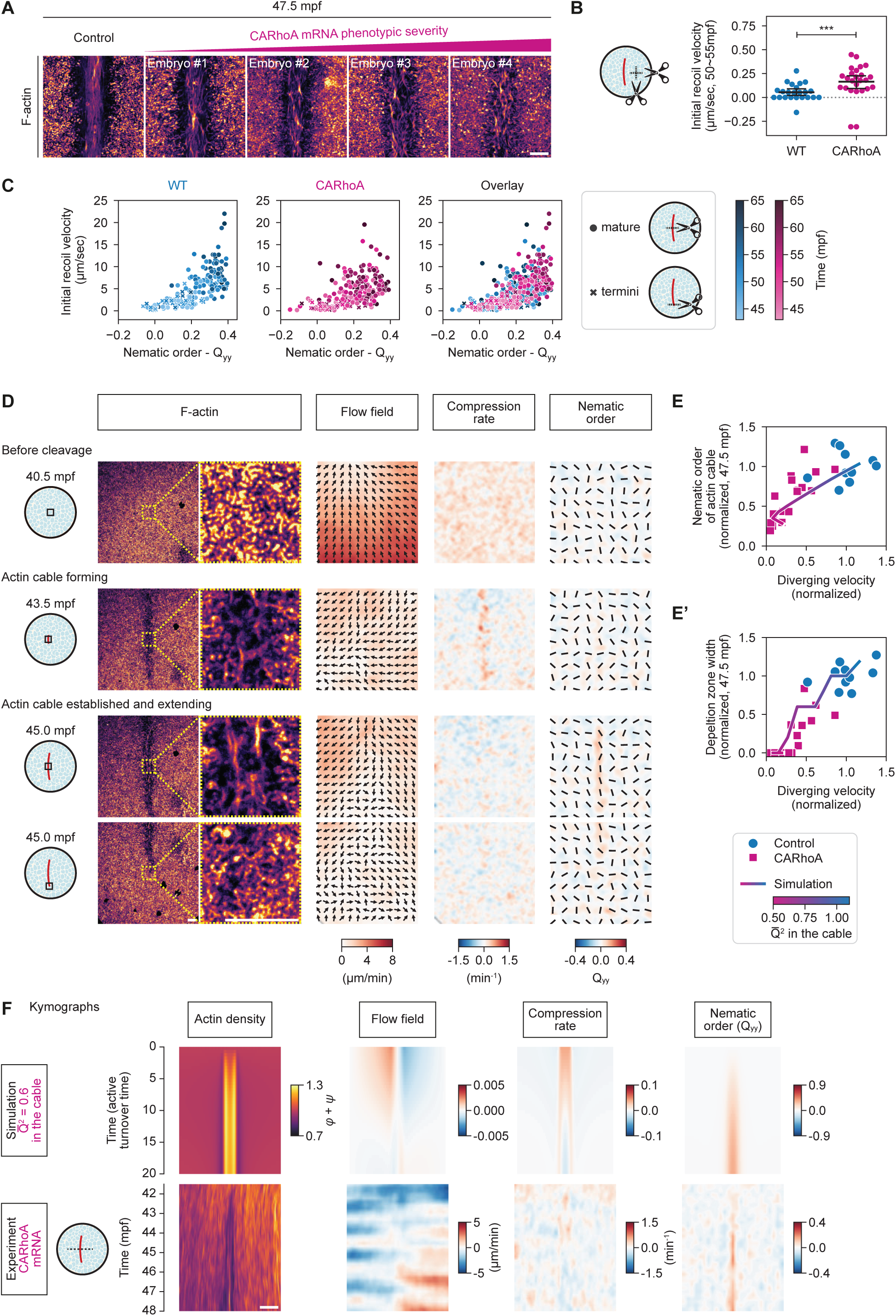
Active gel modelling of changes in cortical actin dynamics upon CARhoA overexpression during the 1st cleavage. (A) Fluorescence images of Tg(*actb2:Lifeact-EGFP*) embryos injected with phenol red only (control) or 600pg CARhoA mRNA (magenta) showing labelled F-actin at 47.5 mpf. Four exemplary embryos were selected and shown in the order of increasing phenotypic severity (from left to right). Scale bar, 10 μm. (B) Plot of initial recoil velocities of the actin cortex after UV-laser ablation in wild-type embryos and embryos injected with 600 pg CARhoA mRNA. Data are shown by a categorical scatter plot with the bold horizontal bars representing the means and the 95% confidence intervals. Wild-type: N = 5 clutches from 2 different days, n = 23 cuts on 14 embryos; CARhoA: N = 11 clutches from 6 different days, n = 25 cuts on 21 embryos. Statistical test, Mann-Whitney test: \*\*\**p* < 0.001. (C) Scatter plots of the initial recoil velocities after laser cutting of the actin cable versus the nematic order (Q_yy_) at the position of laser ablation. Cut positions are indicated by circles (on the mature actin cable) and crosses (on the actin cable termini), as shown by the schematics on the right. Color-coding represents time as shown in the color bars. Wild-type, mature actin cable: N = 14 clutches from 5 different days, n = 134 cuts on 87 embryos; wild-type, actin cable termini: N = 12 clutches from 5 different days, n = 49 cuts on 39 embryos; CARhoA, mature actin cable: N = 21 clutches from 8 different days, n = 202 cuts on 110 embryos; CARhoA, actin cable termini: N = 8 clutches from 4 different days, n = 27 cuts on 24 embryos. (D) Fluorescence images of an exemplary Tg(*actb2:Lifeact-EGFP*) embryo injected with 600 pg CARhoA mRNA showing the labelled cortical actin and their zoom-in views (1st column) at three representative time points during the formation of actin cable, with the corresponding flow field (2nd column; arrows indicate the local flow direction and color-coding the local velocity magnitude), compression rate (3rd column; red indicates compression and blue expansion) and nematic order (4th column; bars indicate the orientation of alignment and color-coding the value of Q_yy_ component; red indicates the alignment along y-axis and blue along x-axis). The position and time points are shown by the schematics on the left. Scale bar, 10 μm. (E, E’) Scatter plots of the nematic order in the actin cable (E) and the width of the actin depletion zones (E’) versus the diverging flow velocities in the surrounding cortex at 47.5 mpf in embryos injected with phenol red only (control, blue) or with 600 pg CARhoA mRNA (magenta). The overlaid line plots are predictions from corresponding simulations, where the nematic stabilization parameter *Q*^2^ was adjusted to mimic varied CARhoA expressing levels, as marked by the color bar below. All experimental data were normalized based on the average value of control data. Control: N = 10 single clutches from 3 different days, n = 10 injected embryos; CARhoA: N = 17 single clutches from 10 different days, n = 17 injected embryos. (F) Kymographs of actin density field (1st column), flow field (2nd column; red indicates movements to the right and blue to the left), compression rate (3rd column; red indicates compression and blue divergence) and nematic order (4th column; red indicates the alignment along y-axis and blue along x-axis) along a line perpendicular to the actin cable at the animal pole in simulations (upper row, with *Q*^2^ adjusted to 0.6 to count for the phenotype observed in CARhoA overexpressing embryos) and experimental observations (lower row) for CARhoA overexpressing embryos (as described in D, E,E’). Scale bar, 20 μm.

To assess whether these effects are also captured by our theoretical model, we modified the model by keeping all contractility parameters constant while reducing the maximum attainable filament alignment (see SI Theory Note for details and for discussion of potential molecular mechanisms through which RhoA could limit actin filament alignment). Consistent with our experimental observations, CARhoA overexpression in the model reduced divergent lateral flows, weakened actin depletion zone formation at the cable’s sides (Fig. 5E, 5E′, S5C, S5D), and decreased the recoil velocity following laser ablation (Fig. S5F). Furthermore, by exploiting variability in the CARhoA phenotype, we examined the functional relationship among these quantities. Simulations with varying CARhoA activity levels predicted that both the nematic order within the actin cable and the width of the actin depletion zone scale linearly with the diverging lateral flow velocity - a prediction that was subsequently confirmed by quantitative analysis of our experimental data (Fig. 5E, 5E′, 5F). Together, these findings demonstrate that the ratio of nematic to isotropic contractility - quantified experimentally and captured by our theoretical framework - is sufficient to account for the observed cortical actin rearrangements and flow patterns during meroblastic cleavages.

## Distinct mechanochemical mechanisms drive cleavage furrow indentation and invagination

To assess the extent to which actin reorganization contributes to blastodisc cytokinesis, we examined how changes in actin architecture correlate with underlying cell shape dynamics (Fig. 6A). Given that furrow indentation during Phase 1 was tightly aligned in both space and time with the formation and extension of the actin cable (Fig. 2A, S2A, Movie S6), the initial furrow ingression might arise from the contraction of the actin cable. In contrast, plasma membrane invagination during Phase 2 was accompanied by the accumulation of Cadherin-1 (Cdh1) at the developing septum^16,17,24^, as detected in Tg(*cdh1-mlanYFP*) embryos (Fig. 6B, Movie S15). This observation raises the possibility that cleavage furrow invagination is mediated by cadherin-dependent membrane adhesion, rather than actomyosin contraction alone.

**Figure 6.**
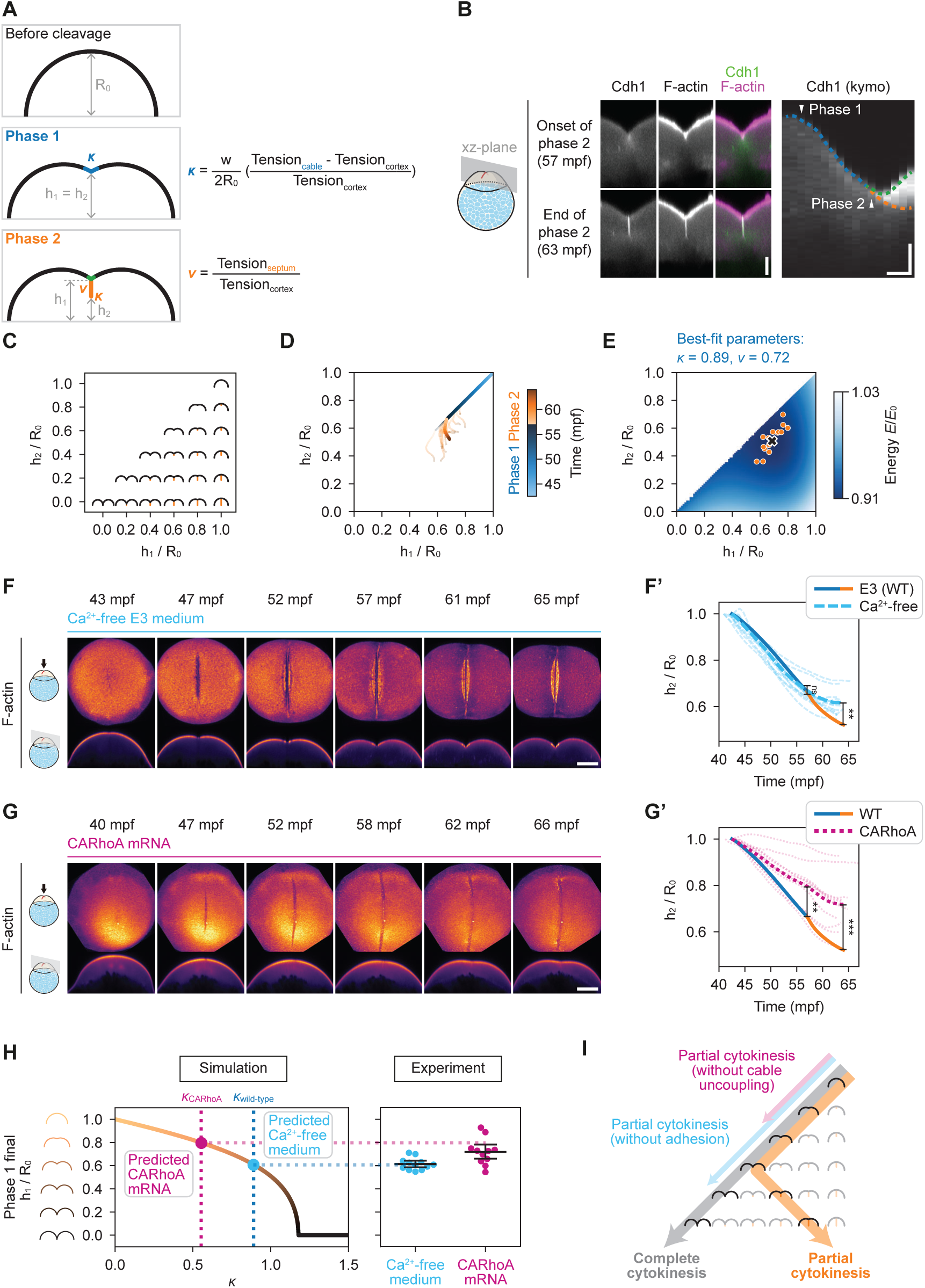
Experimental and theoretical description of cleavage furrow indentation and invagination in wild-type embryos and embryos with compromised cell adhesion and actin dynamics during the 1st cleavage. (A) Schematics of the constriction model for describing the changes in the geometry of embryo surface during the 1st cleavage division. R_0_ indicates the radius of curvature of the animal surface before the 1st cleavage division. h_1_ indicates the height of the furrow ridge, while h_2_ indicates the height of the membrane invagination front during Phase 2 (note that during Phase 1, h_1_ and h_2_ are equivalent). *κ* and *ν* are the two parameters in the model, determining the rescaled contractility of the furrow and the relative tension of the membrane septum, respectively. (B) Fluorescence images of a Tg(*cdh1-mlanYFP*; *actb2:Utrophin-mCherry*) embryo with labelled Cdh1 and F-actin along the xz-plane at the onset (57 mpf) and end (63mpf) of Phase 2 of the 1st cleavage division (left panels) and kymograph of septum invagination along the z-axis in Tg(*cdh1-mlanYFP*) embryos (right panel). Horizontal scale bar, 5 min and vertical scale bar, 50 μm. (C) Schematics illustrating the geometry of the embryo surface (black) and membrane septum (orange) with varied h_1_ and h_2_ normalized by R_0_. Note that only configurations below the diagonal exist, given that h_2_ can not become larger than h_1_. (D) Line graph of h_1_ and h_2_ normalized by R_0_ as a function of time, color-coded by the right color bar. N = 15 clutches from 8 different days, n = 15 embryos. (E) Plot of the theoretically predicted energy landscape as a function of h_1_ and h_2_, determining the geometrical configuration, with the best-fit parameters of *κ* and *ν*. The energy E/E_0_ is color-coded by the right color bar. The orange points indicate experimental data of the final values of h_1_ and h_2_ normalized by R_0_ as in (D) and the black cross the averaged value that is used to fit the energy minimum point of the model. (F, G) Fluorescence images of Tg(*actb2:Utrophin-mCherry*) embryos cultured in Ca^2+^-free E3 medium (F) or injected with 600 pg *CARhoA* mRNA (G) showing labelled F-actin at consecutive time points during the 1st cleavage division. The different orientations are as indicated on the left and in Fig. 1A, 1A’. The top view is a maximum intensity projection. Scale bar, 150 μm. (F’, G’) Line plots showing h_2_ normalized by R_0_ as a function of time. The dashed light-blue line represents the data of embryos cultured in Ca^2+^-free E3 medium (F’), the dotted magenta line embryos injected with 600 pg CARhoA mRNA (G’) and the bold line wild-type embryos cultured in normal E3 medium (F’, G’). Ca^2+^-free: N = 9 clutches from 4 different days, n = 12 embryos; CARhoA: N = 11 clutches from 5 different days, n = 11 embryos. Statistical tests for comparing wild-type and Ca^2+^-free (F’), Mann-Whitney test: at 57 mpf, *p* = 0.981; at the final time point, \*\**p* = 0.007. Statistical tests for comparing wild-type and CARhoA (G’), Mann-Whitney test: at 57 mpf, \*\**p* = 0.002; at the final time point, \*\*\**p* < 0.001 (H) Line graph of simulated h_1_ (normalized by R_0_) at minimum energy, as a function of the parameter *κ* (left panel); on the right are scatter plots, with the bold horizontal bars representing the means and the 95% confidence intervals, of the corresponding experimental data of the final h_1_ (normalized by R_0_) for embryos cultured in Ca^2+^-free medium and embryos injected with 600 pg CARhoA mRNA (right panel; Ca^2+^-free: N = 9 clutches from 4 different days, n = 12 embryos; CARhoA: N = 11 clutches from 5 different days, n = 11 embryos.) In the left panel, vertical dotted lines indicate the *κ* of wild-type and CARhoA data, and the red and blue dots demarcate the predicated final h_1_/R_0_ of embryos cultured in Ca^2+^-free medium (light-blue) and embryos injected with 600 pg CARhoA mRNA (magenta). (I) Progression of shape changes leading to complete/holoblastic cytokinesis (grey) or partial/meroblastic cytokinesis (orange) in schematics illustrating the geometry of the embryo surface (black) and membrane septum (orange) with varied h_1_ and h_2_ as shown in (C). Aberrant shape changes after Ca^2+^-free medium treatment or CARhoA overexpression are shown in light blue and magenta, respectively.

To conceptualize how the combined actions of actin cable contraction and Cdh1-mediated membrane adhesion drive blastodisc cytokinesis, we developed a minimal mathematical model to analyze the coupled mechanics of actin cable contractility, cortical tension, and membrane adhesion during this process (Fig. 6A, see SI Theory Note). By formulating the total elastic potential energy of the actin cortex and contractile cable^9^, we determined the geometric configurations that minimize this energy, allowing us to predict the extent of furrow ingression and invagination resulting from cable contraction and membrane adhesion.

Our model revealed that during Phase 1, when furrow indentation requires deformation of the polar regions, increasing the rescaled contractility (*κ*) of the furrow induces a continuous phase transition from incomplete to complete furrow indentation (see SI Theory Note). This contrasts with classical cytokinetic geometries involving a ring-like actomyosin cable, which predicts a sharp, discontinuous transition at a lower contractility threshold^9^. Crucially, the threshold for continuous furrow indentation depends not only on contractility but also inversely scales with the initial cell radius (Fig. 6A). This implies that in large embryos such as zebrafish, unrealistically high contractility values (10–100 times greater than in typical culture cells) would be required to achieve the full extent of cytokinesis with Phase 1 alone (see SI Theory Note).

To circumvent the need for such extreme contractility, and following our experimental observation of Cdh1 accumulating at the forming septum during Phase 2 (Fig. 6B), we extended our model by incorporating membrane adhesion as an additional mechanism for furrow invagination during Phase 2. In particular, we modelled, given the uncoupling between cable and cortex discussed above, an additional degree of freedom in the system, i.e. that the cable could continue invaginating without cortex deformation, creating a contact zone with relative tension *ν*, which we expect to depend on Cdh1-mediated adhesion^62–64^. With this addition, our model predicts that cytokinesis can proceed substantially further (see SI Theory Note). Moreover, the model reproduces several non-trivial experimental observations, in particular, that in Phase 2, the energetically-favorable path is to decrease furrow depth through invaginating the membrane septum, while moving the contact point slightly upwards (Fig. 6C, 6D, 6E; see SI Theory Note for details). Together, these modelling results predict that successful meroblastic cleavage in large embryos relies on a dual mechanism: actomyosin-driven furrow indentation during Phase 1 and adhesion-mediated membrane invagination during Phase 2 (Fig. 6I).

To experimentally test the predictions of our model - particularly the role of Cdh1-mediated membrane adhesion in furrow invagination - we reduced membrane adhesion by depleting Ca²⁺ from the embryo culture medium, thereby impairing Cdh1 function^65,66^ (Fig. 6F, Movie S16). Strikingly, embryos cultured under Ca²⁺-free conditions exhibited slightly enhanced furrow indentation, yet failed to undergo furrow invagination, ultimately resulting in incomplete blastodisc cytokinesis (Fig. 6F, 6F’, Movie S16). These observations could be qualitatively, but also quantitatively, predicted by our model using the same contractility values as in wild-type, and simply removing the adhesion term: in the absence of membrane adhesion, furrow indentation slightly increases, while membrane invagination is abolished, leading to a net decrease in the extent of cytokinesis (Fig. 6F, 6F’, 6H, S6B).

Finally, we investigated how the formation of actin depletion zones at the lateral sides of the contractile cable influences cytokinesis. To test this, we went back to CARhoA-overexpressing embryos, which exhibited variable degrees of actin depletion zone formation (Fig. 5A). In agreement with the idea that actin depletion zones allowed for furrow invagination, embryos with poor depletion zone formation displayed reduced furrow invagination (Fig. 6G, 6G’, S6B, Movie S17). More quantitatively, simply using the reduced tensile force estimated from the nematic order of actin cable (39% reduction; Fig. S5B) in CARhoA-overexpressing embryos, we could quantitatively predict without free parameters the diminishment of furrow ingression seen in CARhoA-overexpressing embryos (Phase 1 alone; Fig. 6H). Together, these results support the conclusion that successful blastodisc cytokinesis requires the coordinated action of nematic actomyosin cable contraction, actin depletion zones and Cdh1-mediated membrane adhesion (Fig. 6I).

## Discussion

Our findings demonstrate that meroblastic cleavage is driven by the coordinated action of actomyosin cable contraction and Cdh1-mediated membrane adhesion (Fig. 6I). In contrast to the traditional model of cytokinesis - which emphasizes actomyosin ring constriction as the primary force for cell division - our data indicate that, in large cells undergoing partial cytokinesis, the contractile forces generated at the forming cleavage furrow are insufficient to fully separate the animal cytoplasmic region (blastodisc) of the zygote. To overcome this barrier, adhesion between opposing membranes at the tip of the cleavage furrow, and the resulting formation of a membrane fold or septum, are essential for promoting furrow invagination and completing blastodisc separation^67–70^. Membrane adhesion facilitates this process by reducing interfacial tension, thereby enabling expansion of the septum. When combined with actomyosin cable contraction, this promotes efficient and sustained furrow progression.

Crucially, we also show that mechanical uncoupling of the actin cable from the surrounding actin cortex at its lateral edges is necessary for septum invagination. This uncoupling is achieved through an active nematic instability where initial short-range convergent flows lead to nematic alignment, which drastically modifies the geometry of flows towards long-range divergent flows. This eventually leads to the formation of localized actin depletion zones flanking the cable - an architectural feature not observed in conventional cytokinesis. This mechanism may represent a unique adaptation for partial cytokinesis in large cells, where traditional actomyosin-based furrow ingression is mechanically constrained. In line with this, our large-scale model of meroblastic cleavage predicts that this adhesion-based mechanism is beneficial in large embryos but counterproductive in small ones, where it may actually reduce the total extent of furrow indentation and invagination (see Theory Note for details).

Additional cellular features specific to the first meroblastic cleavages in zebrafish may also contribute to cytokinesis. Notably, both our findings and previous studies have identified actin enrichment in protrusions that emerge at the center of the forming cleavage furrow during the late stage of Phase 1^24,71,72^ (Fig. 2B, Movie S5, S7), as well as non-spindle microtubule accumulations near the cleavage site during the transition from Phase 1 to Phase 2^16,18,22^ (Fig. S1A, upper row). It is conceivable that these actin-rich protrusions facilitate the transition between phases by promoting Cdh1-mediated plasma membrane contact at the forming septum. Similarly, the microtubule accumulations may help guide furrow invagination during Phase 2. However, experimentally dissecting the roles of these actin- and microtubule-based structures remains challenging, as selectively targeting them without disrupting the mitotic spindle or cortical actin is technically demanding.

Cytokinesis during early development has largely been studied in embryos undergoing either complete cleavage, as seen in mammals, or superficial cleavage, as observed in insects^73,74^. The mechanisms driving these two cleavage types differ fundamentally: complete cleavages are mediated by actomyosin ring constriction, whereas superficial cleavages occur through membrane invagination in a process commonly referred to as ‘cellularization’. Our study, focusing on meroblastic cleavage in zebrafish, presents a unified model that integrates both processes. We demonstrate how actomyosin ring or cable constriction, in conjunction with membrane adhesion and invagination, can work synergistically to drive cytokinesis. This model provides new insight into the core mechanisms underlying multicellularity and offers a new perspective on its emergence during metazoan evolution.

## Supporting information

SI Theory Note

## Materials and methods

### Zebrafish handling

Zebrafish (*Danio rerio*) maintenance and handling was performed as described in ref.^75^. The following zebrafish strains were used in this study: Tg(*actb2:mCherry-CAAX*)^76^, Tg(*XlEef1a1:dclk2-GFP*)^77^, Tg (*actb2:RFP-pcna*)^78^, Tg(*actb2:Utrophin-mCherry*)^79^, Tg(*actb2:Lifeact-EGFP*)^61^, Tg(*actb2:anillin-mNeonGreen*), Tg(*actb2:Myl12.1-EGFP*)^63^ and Tg(*cdh1-mlanYFP*)xt17^80^. Eggs laid within a period of two minutes were collected. Embryonic manipulations of wild-type embryos were performed in E3 medium (4.96 mM NaCl, 0.18 mM KCl, 0.33 mM CaCl_2_, 0.40 mM MgCl_2_). For the Cdh1-depletion experiment, embryos were collected in normal E3 solution and rinsed in Ca^2+^-free medium (4.96 mM NaCl, 0.18 mM KCl, 0.40 mM MgCl_2_, 0.3 mM EGTA) five times before further manipulation and imaging. All animal experiments were carried out following the guidelines of the Ethics and Animal Welfare Committee (ETK) in Austria.

### Embryo staging

Embryos were staged based on the structural organization of labelled cortical actin, spindle microtubules, cell cycle marker (RFP-PCNA) and their overall geometry (Fig. S1A). The developmental timelines of multiple embryos imaged at 28°C were aligned and averaged to obtain the exact values: onset of mitosis (1st cell cycle) = 30 mpf; onset of interphase (2nd cell cycle) = 40 mpf; onset of 1st cytokinesis (onset of Phase 1) = 42.5 mpf; onset of mitosis (2nd cell cycle) = 55 mpf; onset of interphase (3rd cell cycle) = 64 mpf. Notably, during the rapid cleavage stage of early embryogenesis, the telophase of the 1st cell cycle - when cytokinesis (Phase 1) initiates - overlaps with the interphase of the 2nd cell cycle.

### Molecular biology

To synthesize messenger RNA (mRNA) for the active RhoA sensor mNeonGreen-2xrGBD^36^, the coding sequence was amplified from the plasmid Addgene#129624 with the primers: Fwd 5’-tttgcaggatcccatcgattgccaccATGGTGAGCAAGGGCGAGGA-3’ and Rev 5’-gtaatacgactcactatagttgattatgatcagTTATCTAGAG-3’ and the PCR product was inserted into a pCS2plus plasmid linearized by restriction enzymes BstBI and XbaI using Gibson Assembly Cloning Kit (New England Biolabs) following the manufacturer instructions. The mRNAs for embryo microinjections were synthesized using the mMESSAGE mMACHINE™ SP6 Transcription Kit following the manufacturer instructions.

### Embryo microinjections

Zebrafish embryos were injected using glass capillary needles (30-0020, Harvard Apparatus), which were pulled by a needle puller (P-97, Sutter Instrument) and attached to a microinjector system (PV820, World Precision Instruments). Microinjections of mRNAs were performed at around 8 mpf and the following mRNAs were used: 600 pg *CARhoA*^81^ and 1 ng *mNeonGreen-2xrGBD* (this study). The mRNA was coinjected with a red dye, phenol red. The embryos with the most phenol red in the blastodisc were selected for further experiments.

### Transgenic line generation

The Tg(*actb2:anillin-mNeonGreen*) line was generated in this study. The coding sequence of *anln* was amplified from a cDNA library of larvae stage wild-type AB embryos with the primers: Fwd 5’-ggggacaagtttgtacaaaaaagcaggcttaATGGATCCATTCACCGAGAAACTG-3’ and Rev 5’-ggggaccactttgtacaagaaagctgggtaCATGGGTCTGTAGCAGGAATCC-3’. The PCR product was recombined with *pDONR221* (Lawson#208) and the resulting entry clone was recombined with *pDestTol2pA2* (Chien#394), *p5E β-actin promoter* (Chien#229), *p3E mNeonGreen*^82^ to create *pTol2-actb2:anillin-mNeonGreen*. The *pTol2* vector was co-injected with mRNA encoding the transposase (Invitrogen) into 1 cell-stage wild-type AB embryos. Individual positive carriers were selected and out-crossed with wild-type AB fish for stable single-copy genetic integration.

### Image acquisition

Low-magnification imaging with upright mounting was performed on a Zeiss 900 upright confocal microscope equipped with a Plan-Apochromat 20x/NA 1.0 water dipping objective to characterize embryo geometry and to identify protein localization. Dechorionated embryos were placed in agarose molds and oriented using forceps with the animal pole towards the objective. No agarose or other polymers were placed on top, ensuring that the embryos were neither immobilized nor constrained. High-magnification imaging with inverted mounting was carried out on a Nikon Ti2E inverted microscope equipped with Yokogawa CSU-W1 dual-disk spinning disk unit (50 μm pinholes) and a CFI Plan-Apochromat Lambda 60x/NA 1.4 oil immersion objective to analyze actin organization. Laser ablation assays were performed on the same setup using a CFI Apochromat Lambda 40x/NA 1.15 water immersion objective. For both purposes, dechorionated embryos were mounted in 1% low-melting point agarose (Invitrogen) on glass-bottom dishes (MatTek) and carefully oriented using forceps with the animal pole towards the coverslip. The temperature was set to 28°C for upright imaging and 25°C for inverted imaging.

### Laser ablation assay

Laser ablation assays were carried out with a custom-built setup. Live samples were ablated along 15 μm-long lines using a Teem Photonics UV (355 nm) pulsed laser (13 μJ per pulse at 5 kHz) applied to 30 equidistant sites (25 pulses per site) and imaged at exposure times of 200 milliseconds. To determine initial recoil velocities, a line was drawn perpendicular to the ablation site to generate a kymograph, from which the opening distances after ablation were extracted either manually or with a custom Python script. A linear fit of distance (μm) versus time (second) was performed from the first post-ablation frame until the interval showing the best linear regression (typically within 10 frames), with the slope taken as the initial recoil velocity. Cases showing a wound response after ablation were excluded from analysis. For the long-cut experiment, embryos were ablated along an 80 to 120 μm-long line applied to 200 equidistant sites (25 pulses per site).

### Data analysis and statistics

All image data were analyzed using Fiji (National Institutes of Health)^83^ and custom Python scripts incorporating NumPy, SciPy, scikit-Image, and scikit-learn. Data handling and visualization were performed with Microsoft Excel, Adobe Illustrator and custom Python scripts with NumPy, SciPy, statsmodel^84^, Pandas, Matplotlib and Seaborn.

#### Surface curvature quantification and cleavage furrow detection

As illustrated in Fig. S1B, time-lapse z-stack movies of embryos with labelled plasma membrane were segmented using a custom Python script to extract local surface heights in the form *z* = *f*(*x, y*). Mean curvature (*H*) and Gaussian curvature (*K*) were computed from the first and second derivatives of the surface using standard expressions from differential geometry. The principal curvatures (*k*_1_ and *k*_2_) were then obtained from these quantities 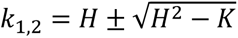. The cleavage furrow was identified as a region exhibiting positive values of the principal curvature *k*_1_. The membrane invagination front during Phase 2 was manually marked in Fiji and the 3D coordinates were exported and further processed in a custom Python script for analysis and illustration.

#### Quantification of furrow indentation and membrane invagination

The positions of the furrow ridge and the membrane invagination front along the z-axis were manually marked on the central xz-plane perpendicular to the cleavage furrow. Their z-axis coordinates were referenced to the onset of cleavage, yielding negative displacement values (Fig. 1F, 1F’). To normalize these displacements, R_0_, h_1_ and h_2_ were measured as defined in Fig. 6A. Points at the center of the animal surface were selected and fit to a circle to determine the local radius of curvature, taken as R_0_. The negative z-axis coordinates of the furrow ridge and the membrane invagination front were then added to R_0_ to obtain h_1_ and h_2_, respectively.

#### Flow field and compression rate

The flow field was analyzed with particle image velocimetry (PIV) using PIVlab running on Matlab^85^. Time-lapse movies acquired at 5-10 second intervals were processed using two successive passes through interrogation windows of 64/32. To remove noises and invalid vectors, both a standard deviation filter and a local median filter were applied, and the few missing vectors were subsequently interpolated. The resulting velocity fields were exported to a custom Python script to calculate the compression rate, defined as negative divergence, −*∂_x_v_x_ − ∂_y_v_y_*.

#### Nematic order quantification

The nematic order parameters were quantified using an indirect method based on Fourier Transform analysis of fluorescence image intensity as described in ref.^47^. The size of square regions was 40 pixels (6.46 μm) and *q*_min_ and *q*_max_ were set to be 2 and 20, respectively.

*Quantification of nematic order of the actin cable after UV-laser ablation (in Fig. S3B, S3B’)*

The analysis was performed on datasets in which the actin cable remained within the focal plane following the UV-laser ablation on it. The nematic order within a square region of 40 pixels (6.46 μm) centered on the ablation site was quantified and plotted as a function of time. For image visualization, due to the pronounced drop of actin intensity after ablation, the raw images were adjusted using gamma correction (*γ* = 0.35) for better display.

#### Quantification of the depletion zone width

The width of the regions with actin intensity 25% lower than that in the actin cable was manually measured for experimental data, while for simulations, the width of the regions where the actin density was 0.1 lower than that in the actin cable was measured.

#### Quantification of furrow ridge angle

The cleavage furrow was outlined on the central xz-plane (perpendicular to the furrow) using a custom script. Tangent lines were fitted to the surface on either side of the furrow ridge, and the furrow ridge angle was defined as half of the angle between these tangents.

#### Statistics

The statistical analyses were carried out using statistical functions in SciPy and statsmodel^84^. The number of eggs (n) and experimental replicates (N; defined as clutches laid within a two-minute period, with at least a one-hour interval between successive clutches) are indicated in the figure legends. All experiments were repeated independently at least three times. Error bars in all figures represent the 95% confidence interval.

## Acknowledgements

We are grateful to the members of the Hannezo and Heisenberg groups for discussions and technical advice. We also thank the Imaging and Optics Facility and the Lab Support Facility at ISTA for their continuous support. Y.I.L. acknowledges funding from the European Union’s Horizon 2020 research and innovation programme under the Marie Skodowska-Curie Grant Agreement No. 101034413. The research was supported by funding to C.-P.H. from the NOMIS Foundation (Project ID 1.844) and to E.H. from the European Research Council (ERC) under the European Union’s Horizon 2020 research and innovation programme (grant agreement no. 851288).

## Author contributions

X.T., Y.I.L., E.H. and C.-P.H. conceptualized the project, acquired funding, designed experiments and wrote the manuscript. X.T. and J.S. performed experiments. X.T. and Y.I.L. analysed the data. Y.I.L. and E.H. wrote the theory and performed numerical simulations.

---

**Figure S1.**
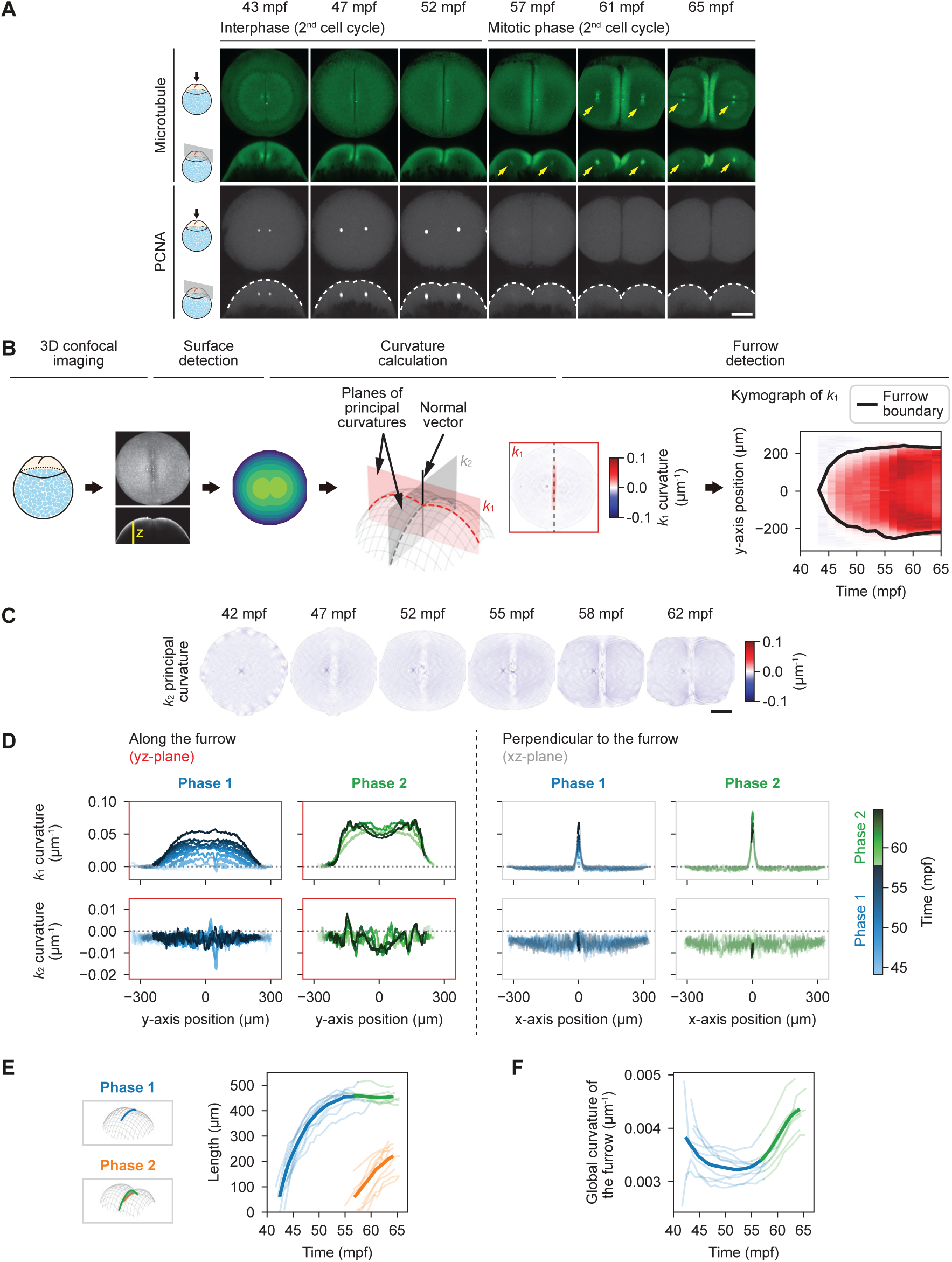
Developmental staging of cleavage furrow geometry during the 1st cleavage, related to Figure 1. (A) Fluorescence images of a Tg(*XlEef1a1:dclk2-GFP*) embryo with labelled microtubules (upper row) and a Tg(*actb2:RFP-pcna*) embryo with labelled nuclei (lower row) at consecutive times during the 1st cleavage division. The different orientations are as indicated on the left and in Fig. 1A, 1A’. The top and side views are maximum intensity projections. Yellow arrows indicate the mitotic spindle during the mitotic phase and white dashed lines the outline of the animal surface. Scale bar, 150 μm. (B) Workflow of the geometry analysis. The confocal fluorescence images of Tg(*actb2:mCherry-CAAX*) embryos were segmented to obtain the surface height map. The principal curvatures of the animal surfaces were calculated. The *k*_1_ and *k*_2_ components indicate the local curvatures along the xz-plane (the red plane) and the yz-plane (the grey plane), respectively. A kymograph of *k*_1_ curvature was made along the cleavage furrow, and the *k*_1_ curvature value was thresholded to determine the two termini of the cleavage furrow. (C) *k*_2_ component of the principal curvature of the animal surface at consecutive time points during the 1st cleavage division. Scale bar, 150 μm. (D) Line plots showing the distributions of *k*_1_ and *k*_2_ principal curvatures on the animal surface along the yz-plane (red frame) and xz-plane (grey frame) at consecutive time points during Phase 1 and Phase 2, color-coded as shown in the left color bar. The upper row panels are along the furrow and the lower row panel perpendicular to it. The bold part of the line indicates the region with positive *k*_1_ curvature (the position of the geometrical furrow ridge). (E) Line graph showing the lengths of the furrow ridge and the membrane invagination front as a function of time during the 1st cleavage division. Color illustrations are shown on the left and in Fig. 1C. N = 9 clutches from 5 different days, n = 9 embryos. (F) Line graph showing the total curvature of the furrow ridge as a function of time during the 1st cleavage division. Color illustrations are the same as in (E). N = 9 clutches from 5 different days, n = 9 embryos.

**Figure S2.**
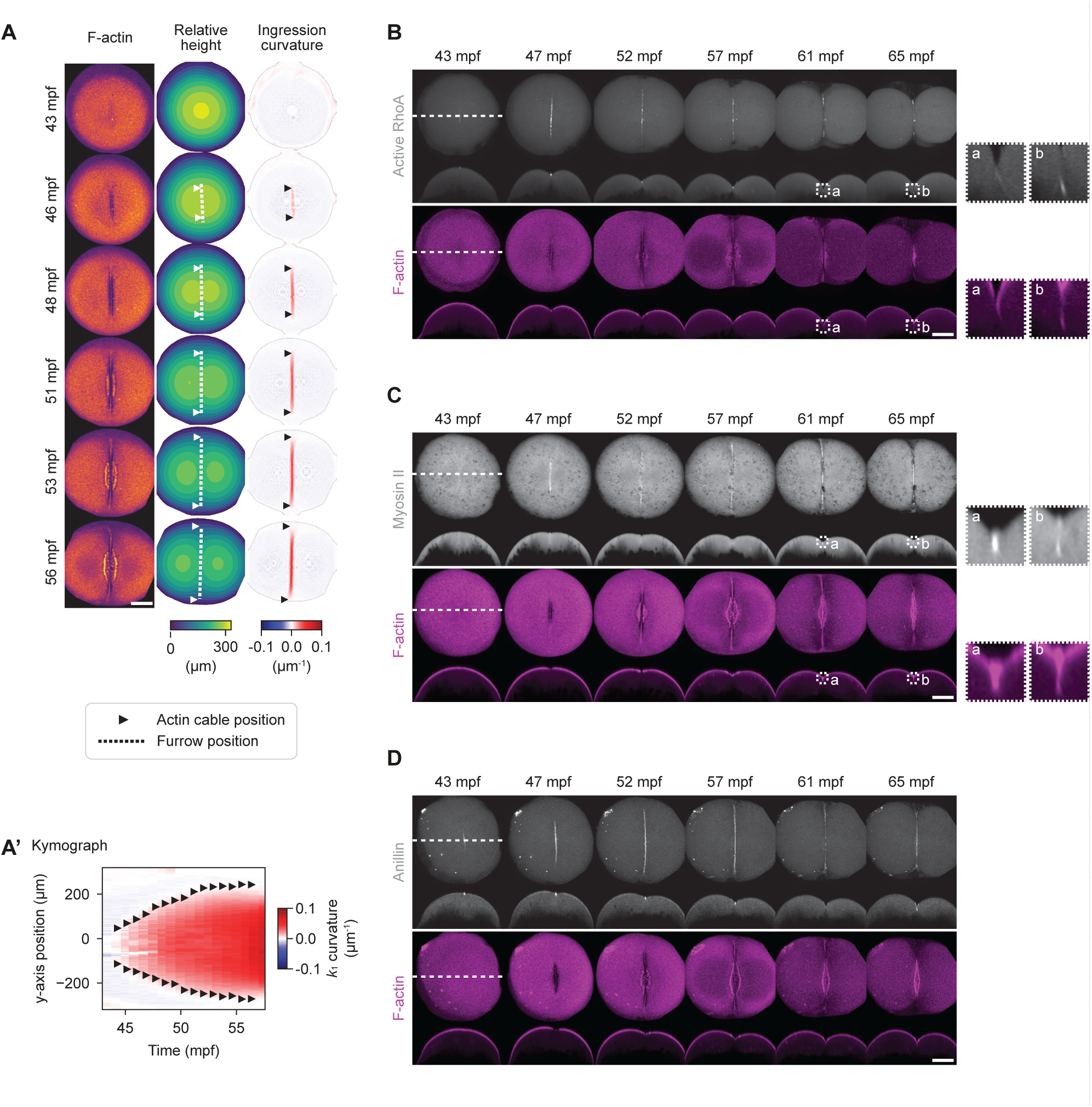
Geometrical and biochemical properties of the forming actin cable during the 1st cleavage, related to Figure 2. (A) Fluorescence images of a Tg(*actb2:Utrophin-mCherry*) embryo with labelled F-actin (left), the surface height map (middle) and the surface ingression curvature (*k*_1_ principal curvature; right) at consecutive time points during Phase 1 of the 1st cleavage division. Arrow heads indicate the two termini of the actin cable, and dotted lines the position of the furrow ridge determined by positive surface ingression curvature. Scale bar, 150 μm. (A’) Kymograph of the ingression curvature along the cleavage furrow (a middle plane along y-axis) during the 1st cleavage division. Arrow heads indicate the positions of the two actin cable termini. (B, C, D) Fluorescence images of a wild-type embryo injected with 1 μg mRNA of mNeonGreen-2xrGBD labelling active RhoA (B), a Tg(*actb2:Myl12.1-EGFP*) embryo with labelled myosin-2 light chain (C), and a Tg(*actb2:anillin-mNeonGreen*) embryo with labelled Anillin (D) at consecutive time points during the 1st cleavage division with top and sectional views (shown in Fig 1A, 1A’). Top views are maximum intensity projections and sectional views are single planes along the dotted lines. Scale bar, 150 μm.

**Figure S3.**
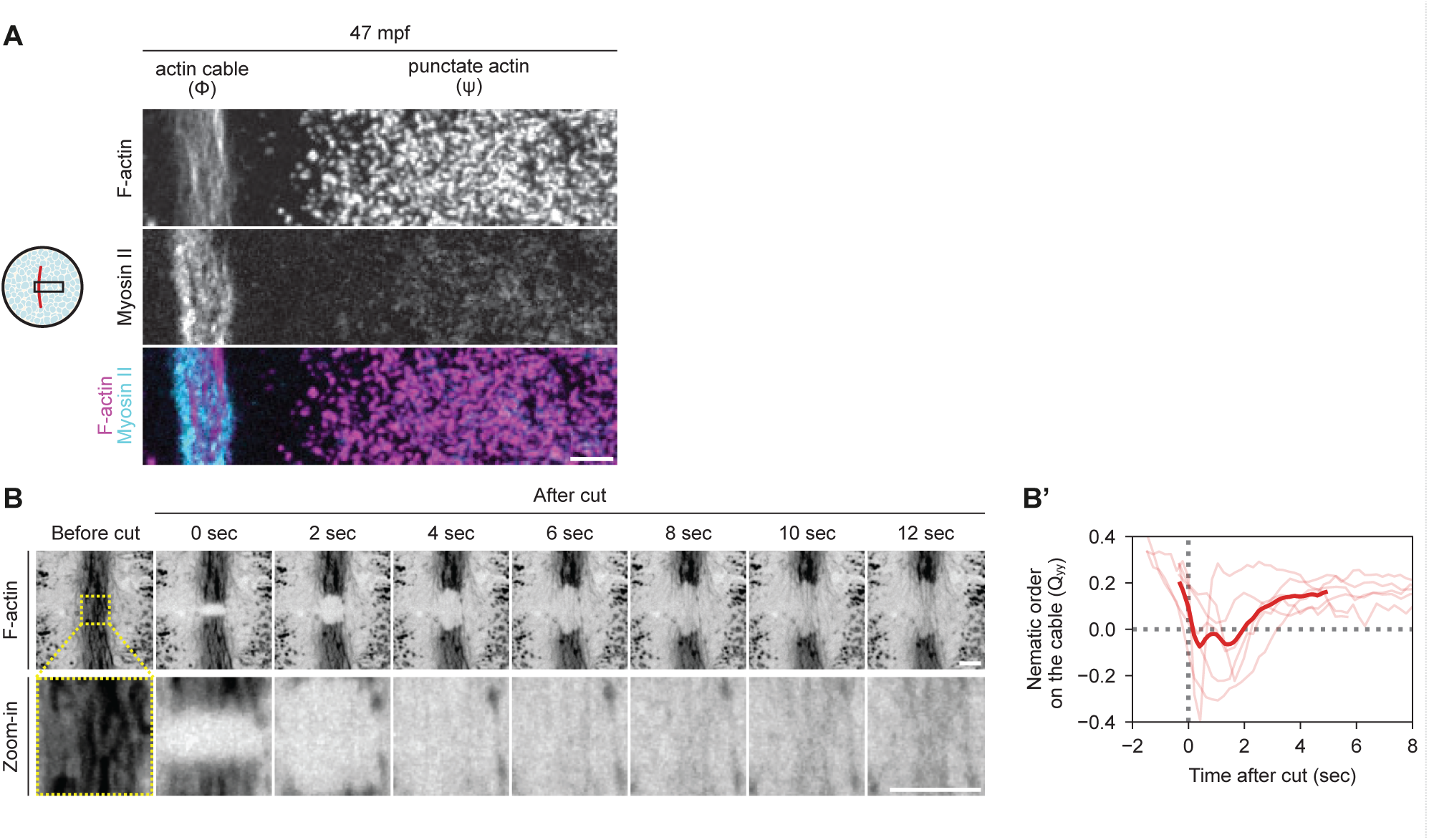
Myosin-2 localization in the actin cable and adjacent cortex and nematic alignment recovery after UV-laser ablation during the 1st cleavage, related to Figure 3. (A) Fluorescence images of a Tg(*acbt2:Utrophin-mCherry*; *actb2:myl12.1-EGFP*) embryo with labelled F-actin and myosin-2 light chain, showing their localizations in the actin cable and surrounding cortex (image position is shown by the schematics on the left) at 47 mpf. Scale bar, 5 μm. (B) Fluorescence images of an exemplary UV-laser ablation of the actin cable in Tg(*actb2:Lifeact-EGFP*) embryos with labelled F-actin at 45 mpf showing the recovery of nematic order at consecutive time points before and after ablation. The zoom-in view, based on which the nematic order was computed, was shown in the lower row. Scale bar, 5 μm. (B’) Line plot of nematic alignment recovery of the actin cable along the y-axis (Q_yy_) as a function of time after UV-laser ablations at 45 mpf. N = 7 clutches from 3 different days, n = 7 cuts on 7 embryos.

**Figure S4.**
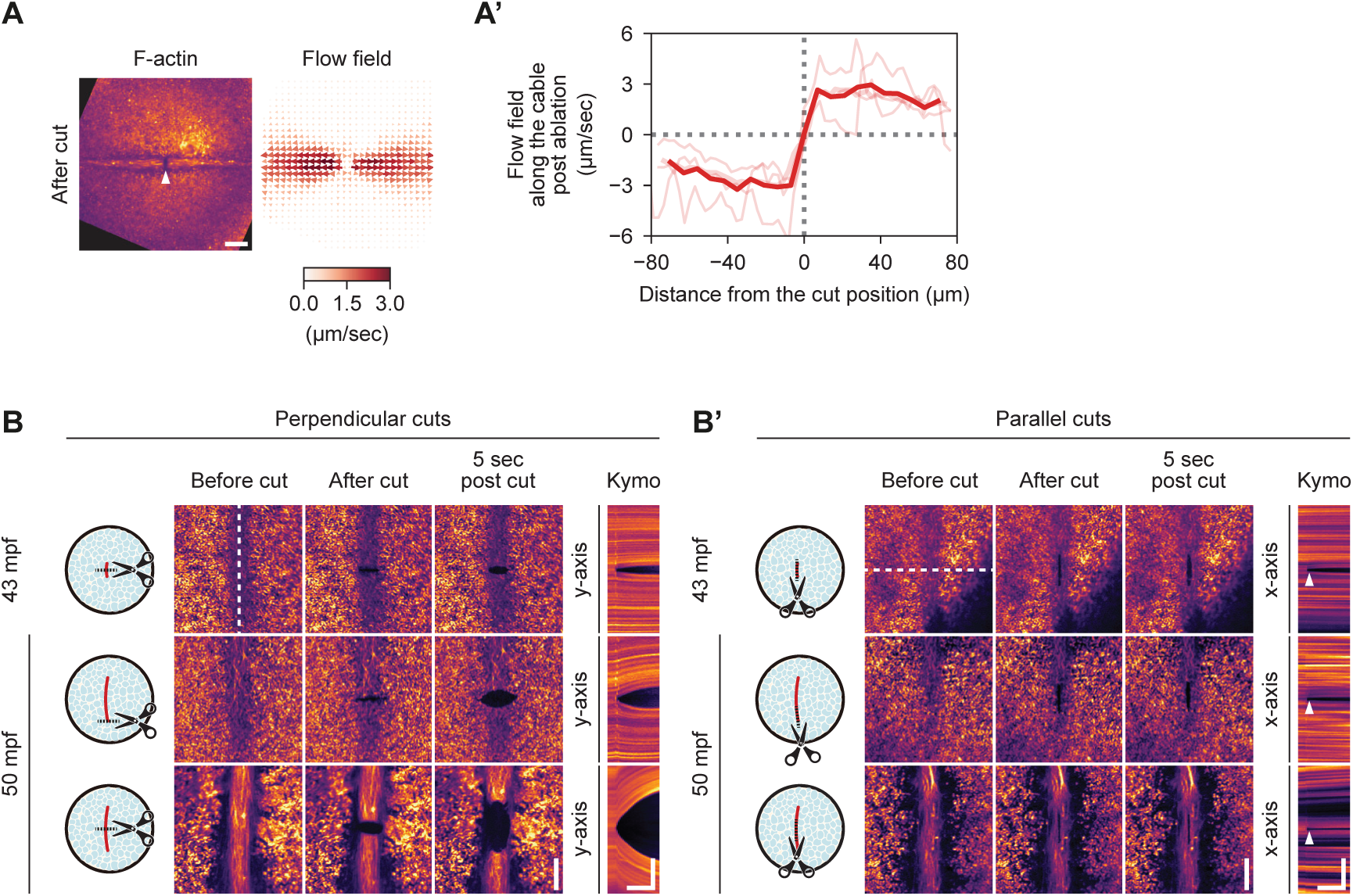
Actin flow dynamics after UV-laser ablation and exemplary images for the ablation experiments shown in Figure 4B, 4B’, related to Figure 4. (A) Fluorescence image of an exemplary UV-laser ablation of the actin cable in a Tg(*actb2:Lifeact-EGFP*) embryo with labelled F-actin (left panel) and the corresponding flow field at the same point (right panel) at 46 mpf. The arrow head on the left panel indicates the position of ablation. The magnitude of local velocities are indicated by both the arrow length and color-coding (as shown in the color bar below). Scale bar, 20 μm. (A’) Line graph of the flow field along the actin cable after UV-laser ablation. Note that the negative values indicate movements toward left and the positive values toward right. N = 4 clutches from 2 different days, n = 4 cuts. (B, B’) Fluorescence images of exemplary UV-laser ablations for the results shown in Fig. 4B, 4B’. The actin cable in Tg(*actb2:Lifeact-EGFP*) embryos with labelled F-actin was cut either perpendicularly (B) or in a parallel manner (B’) at 43 mpf (forming actin cable, upper row) and 50 mpf (mature actin cable, middle and lower row). The cut positions and orientations are shown by the schematics on the left. The fluorescence images before cut, 0 and 5 seconds after cut are shown in the left three columns. Kymographs made along the y-axis (B) or x-axis (B’), indicated by the white dashed lines, are shown in the right column. The arrow heads (B’) indicate the position of the cut. Horizontal scale bar, 5 sec and vertical scale bar, 10 μm.

**Figure S5.**
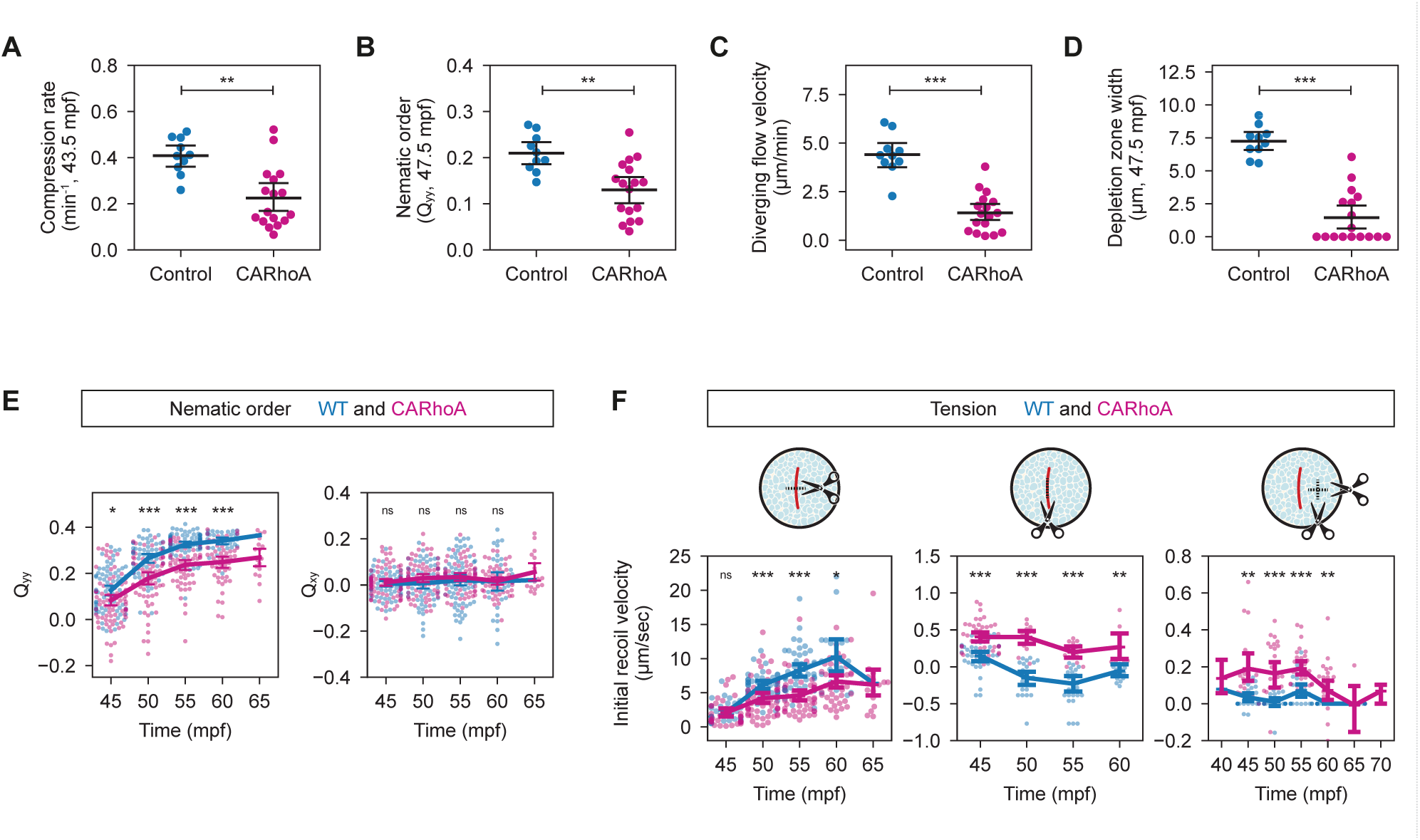
Changes in cortical actin dynamics upon CARhoA overexpression during the 1st cleavage, related to Figure 5. (A, B, C, D) Plots of compression rate of the initial convergent flow at 43.5 mpf (A), nematic order (Q_yy_ component) of the actin cable at 47.5 mpf (B), the diverging flow velocity (C) and the widths of actin depletion zones at 47.5 mpf in embryos injected with phenol red only (control) and 600 pg CARhoA mRNA. Data are shown in categorical scatter plots with the bold horizontal bars representing the means and the 95% confidence intervals. Control: N = 10 single clutches from 3 different days, n = 10 injected embryos; CARhoA: N = 17 single clutches from 10 different days, n = 17 injected embryos. Statistical tests: (A) Mann-Whitney test, \*\**p* = 0.002; (B) Student’s t-test, \*\**p* = 0.001; (C) Student’s t-test, \*\*\**p* < 0.001; (D) Mann-Whitney test, \*\*\**p* < 0.001. (E) Line plots of the nematic order in the actin cable (Q_yy_ component, left panel and Q_xy_, right panel) as a function of time during the 1st cleavage division. Wild-type: N = 21 clutches from 8 different days, n = 145 embryos; CARhoA: N = 22 clutches from 8 different days, n = 131 embryos. Statistical tests, Mann-Whitney tests to compare wild-type and CARhoA data at each time point, with P-values adjusted for multiple comparisons using the Bonferroni correction: Q_yy_, from 45 to 60 mpf, \**p* = 0.043, \*\*\**p* < 0.001, \*\*\**p* < 0.001, \*\*\**p* < 0.001; Q_xy_, from 45 to 60 mpf, *p* = 1, *p* = 0.977, *p* =1, *p* = 1. (F) Line plots of the initial recoil velocities after laser cutting at different positions as shown in the corresponding schematics above as a function of time during the 1st cleavage division. Perpendicular cuts on the cable: wild-type, N = 14 clutches from 5 different days, n = 134 cuts on 87 embryos, CARhoA, N = 21 clutches from 8 different days, n = 202 cuts on 110 embryos; parallel cuts on the cable: wild-type, N = 10 clutches from 5 different days, n = 102 cuts on 59 embryos, CARhoA, N = 16 clutches from 7 different days, n = 74 cuts on 58 embryos; cuts on the surrounding cortex: wild-type, N = 5 clutches from 2 different days, n = 104 cuts on 48 embryos, CARhoA, N = 14 clutches from 7 different days, n = 116 cuts on 67 embryos. Statistical tests, Mann-Whitney tests to compare wild-type and CARhoA data at each time point, with P-values adjusted for multiple comparisons using the Bonferroni correction: perpendicular on the actin cable, from 45 to 60 mpf, *p* = 1, \*\*\**p* < 0.001, \*\*\**p* < 0.001, \**p* = 0.026; parallel on the actin cable, from 45 to 60 mpf, \*\*\**p* < 0.001, \*\*\**p* < 0.001, \*\*\**p* < 0.001, \*\**p* < 0.002; on the surrounding cortex, from 45 to 60 mpf, \*\**p* = 0.001, \*\*\**p* < 0.001, \*\*\**p* < 0.001, \*\**p* = 0.008.

**Figure S6.**
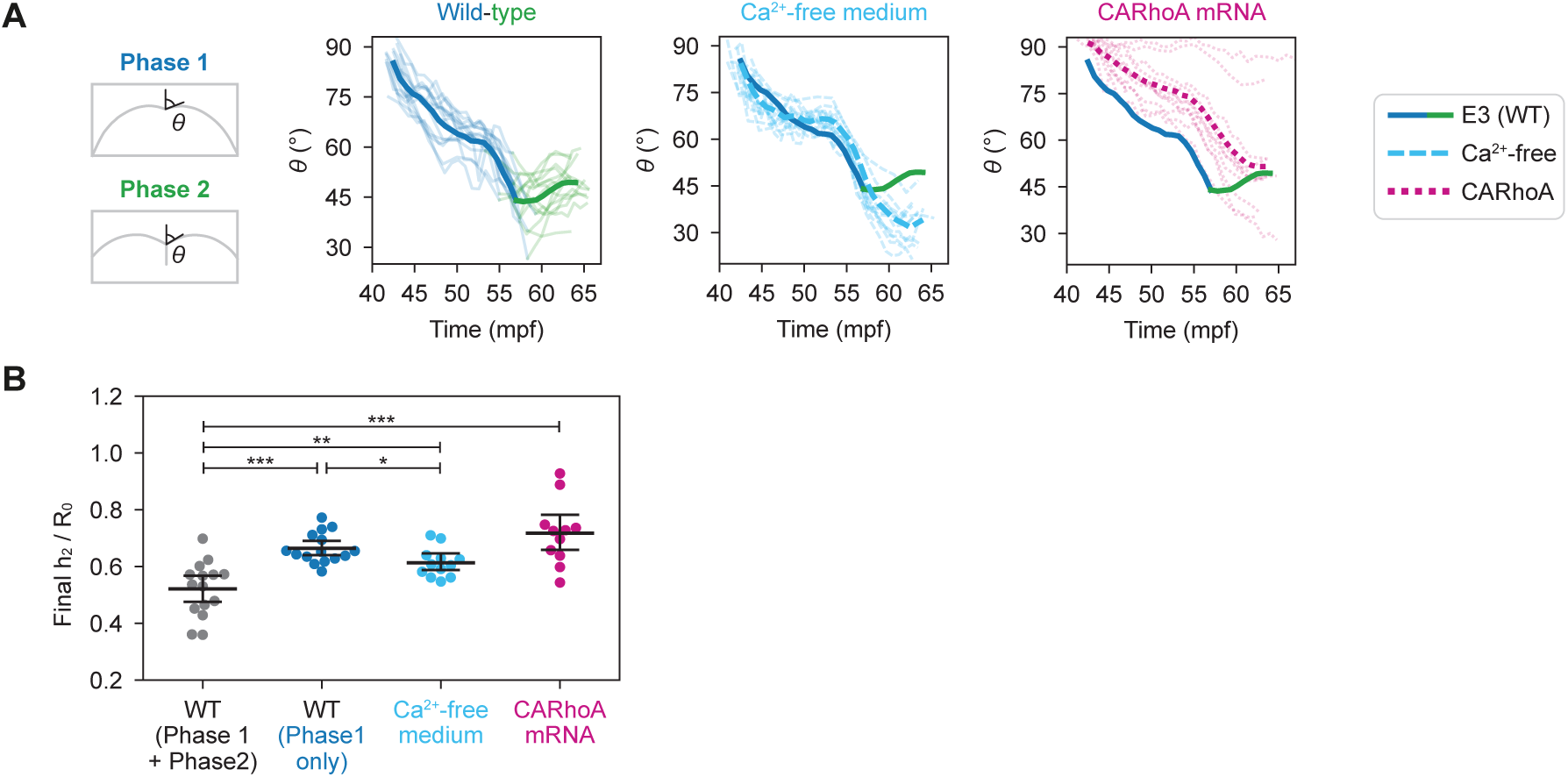
Cleavage furrow angle and indentation in wild-type embryos and embryos with compromised cell adhesion and actin dynamics during the 1st cleavage, related to Figure 6. (A) Line plots of the angle of the furrow ridge (illustrated on the left) as a function of time in wild-type embryos (left panel), embryos cultured in Ca^2+^-free medium (middle panel), and embryos injected with 600 pg CARhoA mRNA (right panel). The bold line colored in blue and green show the data from wild-type, the dashed line colored in light blue the embryos cultured in Ca^2+^-free medium and the dotted line colored in magenta the embryos injected with 600 pg CARhoA mRNA. Wild-type: N = 15 clutches from 8 different days, n = 15 embryos; Ca^2+^-free: N = 9 clutches from 4 different days, n = 12 embryos; CARhoA: N = 11 clutches from 5 different days, n = 11 embryos. (B) Plot of the final h_2_ (normalized by R_0_) at the end of cytokinesis in wild type (gray dots), at the end of Phase 1 in wild type (darker blue), at the end of cytokinesis in embryos cultured in Ca^2+^-free medium (light blue), and at the end of cytokinesis in embryos injected with 600 pg CARhoA mRNA (magenta). Data are shown in a categorical scatter plot with the bold horizontal bars representing the means and the 95% confidence intervals. Wild-type: N = 15 clutches from 8 different days, n = 15 embryos; Ca^2+^-free: N = 9 clutches from 4 different days, n = 12 embryos; CARhoA: N = 11 clutches from 5 different days, n = 11 embryos. Statistical test: Mann-Whitney tests for comparing WT (Phase1 + Phase 2) and Ca^2+^-free, \*\**p* = 0.027, WT (Phase 1 only) and Ca^2+^-free, \**p* = 0.048, WT (Phase 1 + Phase 2) and CARhoA, \*\*\**p* < 0.001; paired Student’s t-test for comparing WT (Phase 1 + Phase 2) and WT (Phase 1 only), \*\*\**p* < 0.001 (P-values were adjusted for multiple comparisons using the Bonferroni correction).

## Movie legends

**Movie S1. The 1st meroblastic cleavage of a zebrafish embryo, related to Fig. 1B**.

Bright-field movie of a dechorionated zebrafish embryo during the 1st cleavage division (39 - 65 mpf, 52 sec/frame). Side view (as indicated in Fig. 1A). Scale bar, 150 μm.

**Movie S2. The 1st cleavage division of a zebrafish embryo, related to Fig. 1B**.

Fluorescence movie of a Tg(*actb2:mCherry-CAAX*) zebrafish embryo with labelled plasma membrane during the 1st cleavage division (40 - 65 mpf, 100 sec/frame). The different orientations are as indicated in the corresponding schematics and in Fig. 1A, 1A’. The top and side views are maximum intensity projections. Scale bar, 100 μm.

**Movie S3. Changes in surface geometry during the 1st cleavage division of a zebrafish embryo, related to Fig. 1D, 1E, S1C, S1D**.

Movie illustrating the surface geometry analysis (40 - 65 mpf, 100 sec/frame). Scale bar, 100 μm.

(Upper row) Top view of a Tg(*actb2:mCherry-CAAX*) zebrafish embryo with labelled plasma membrane (1st panel) was segmented and converted to a height map (2nd panel), based on which the two principal curvatures, *k*_1_ (3rd panel) and *k*_2_ (4th panel), were computed.

(Middle and lower rows) The relative height of the surface (1st panel), the *k*_1_ (2nd panel) and the *k*_2_ curvature (3rd panel) along the yz-sectional plane (middle row; the red bold line in the height map) and xz-sectional plane (lower row; the grey bold line in the height map).

**Movie S4. 3D surface shape changes during the 1st cleavage division of a zebrafish embryo, related to Fig. 1D**.

3D-reconstructed movie of the plasma membrane during the 1st cleavage division (40 - 65 mpf, 100 sec/frame). The wireframe indicates the embryo surface, with the bold blue line marking the cleavage furrow ridge during Phase 1, the bold green line marking the furrow ridge during Phase 2 and the bold orange line marking the membrane contact invagination front, as illustrated in Fig. 1C. The grey shaded area filled between the green and orange lines indicates the membrane contact septum. Scale bar, 100 μm.

**Movie S5. Cortical actin reorganization during the 1st cleavage division of a zebrafish embryo (low magnification), related to Fig. 2B**.

Low-magnification fluorescence movie of a Tg(*actb2:Utrophin-mCherry*) zebrafish embryo with labelled F-actin during the 1st cleavage division (41 - 65 mpf, 52 sec/frame). The different orientations, the top view (upper panel) and the xz-sectional view (lower panel), are as indicated in Fig. 1A, 1A’. The top view is a maximum intensity projection. Scale bar, 150 μm.

**Movie S6. Actin cable formation within the cleavage furrow during the 1st cleavage division of a zebrafish embryo, related to Fig. 2A. S2A.**

Fluorescence movie of a Tg(*actb2:Utrophin-mCherry*) embryo with labelled F-actin (left), the surface height map (middle) and the surface ingression curvature (*k*_1_ principal curvature; right) during Phase 1 of the 1st cleavage division (41 - 57 mpf, 52 sec/frame). Arrow heads indicate the two termini of the actin cable, and dotted lines the position of the furrow ridge determined by positive surface ingression curvature. Color bars are the same as in Fig. 2A. Scale bar, 100 μm.

**Movie S7. Cortical actin reorganization during the 1st cleavage division of a zebrafish embryo (high magnification), related to Fig. 2C**.

High-resolution fluorescence movie of a Tg(*actb2:Lifeact-EGFP*) embryo with maximum intensity projection of a 15 μm z-stack showing the organizations of F-actin during the 1st cleavage division (41 - 50 mpf, 5.4 sec/frame). Overview (left panel) and zoom-in view marked by a white square (right panel). Scale bar, 10 μm.

**Movie S8. Dynamic changes of the flow field, compression rate and nematic order of the actin cortex during the 1st cleavage division of a zebrafish embryo, related to Fig. 2F**.

Fluorescence movie of a Tg(*actb2:Lifeact-EGFP*) embryo with labelled cortical actin during Phase 1 of the 1st cleavage division (1st panel), with the corresponding flow field (2nd panel; arrows indicating the local flow direction and color-coding the local velocity magnitude), compression rate (3rd panel; red indicating compression and blue expansion) and nematic order (4th panel; bars indicate the orientation of alignment and color-coding the value of Q_yy_ component; red indicates the alignment along y-axis and blue along x-axis) during the 1st cleavage division (41 - 50 mpf, 5.4 sec/frame). Scale bar, 20 μm.

**Movie S9. Nematic order recovery on the actin cable after UV-laser ablation during the 1st cleavage division of a zebrafish embryo, related to Fig. S3B.**

Fluorescence movie of an exemplary UV-laser ablation of the actin cable in Tg(*actb2:Lifeact-EGFP*) embryos with labelled F-actin at 45 mpf showing the recovery of nematic order at consecutive time points before and after ablation (0.33 sec/frame). The zoom-in view, based on which the nematic order was computed, was shown in the right panel. Scale bar, 5 μm.

**Movie S10. UV-laser ablation on the actin cable and adjacent cortex during the 1st cleavage division of a zebrafish embryo, related to Fig. 4A**.

Fluorescence movie of UV-laser ablations of the actin cable or adjacent cortex in Tg(*actb2:Lifeact-EGFP*) embryos with labelled F-actin at 50 mpf (0.2 sec/frame). The cut positions and orientations are shown by the corresponding schematics above. Scale bar, 10 μm.

**Movie S11. Dynamics of the actin flow field during and after UV-laser ablation of the actin cable, related to Fig. S4A.**

Fluorescence movie of an exemplary UV-laser ablation of the actin cable in a Tg(*actb2:Lifeact-EGFP*) embryo with labelled F-actin (left panel) and the corresponding flow field at the same point (right panel) at 46 mpf (0.2 sec/frame). The magnitude of local velocities are indicated by both the arrow length and color-coding (as shown in the color bar below). Scale bar, 50 μm.

**Movie S12. Dynamics of the actin flow field during and after a 80 μm-long UV-laser ablation across the actin cable, related to Fig. 4C**.

Fluorescence movie of an exemplary 80 μm-long UV-laser cut perpendicular to the actin cable in a Tg(*actb2:Lifeact-EGFP*) embryo during and after ablation (left panel) and the corresponding flow fields (right panel) at 50 mpf (0.2 sec/frame). The magnitude of local velocities are indicated by both the arrow length and color-coding (as shown in the color bar below). Scale bar, 20 μm.

**Movie S13. Cortical actin reorganization during the 1st cleavage division of a zebrafish embryo overexpressing CARhoA, related to Fig. 5A**.

Fluorescence movies of Tg(*actb2:Lifeact-EGFP*) embryos injected with phenol red only (control) or 600pg CARhoA mRNA (magenta) showing labelled F-actin during the 1st cleavage division (42 - 57 mpf, 10.5 sec/frame). Four exemplary embryos were selected and shown in the order of increasing phenotypic severity (from left to right). Scale bar, 10 μm.

**Movie S14. Dynamic changes of the flow field, compression rate and nematic order of the actin cortex during the 1st cleavage division of a zebrafish embryo overexpressing CARhoA, related to Fig. 5D**.

Fluorescence movie of a Tg(*actb2:Lifeact-EGFP*) embryo injected with 600 pg CARhoA mRNA with labelled cortical actin during Phase 1 of the 1st cleavage division (40 - 50 mpf, 10.5 sec/frame), with the corresponding flow field (2nd panel; arrows indicating the local flow direction and color-coding the local velocity magnitude), compression rate (3rd panel; red indicating compression and blue expansion) and nematic order (4th panel; bars indicate the orientation of alignment and color-coding the value of Q_yy_ component; red indicates the alignment along y-axis and blue along x-axis). Scale bar, 20 μm.

**Movie S15. Dynamic changes in Cdh1 accumulation within the cleavage furrow (septum) during the 1st cleavage division of a zebrafish embryo, related to Fig. 6B**.

Fluorescence movie of a Tg(*cdh1-mlanYFP*; *actb2:Utrophin-mCherry*) embryo with labelled Cdh1 and F-actin during the 1st cleavage of the zygote (39 - 66 mpf, 94 sec/frame). The yellow arrow indicates the membrane septum during Phase 2. The different orientations, the top view (upper row) and the xz-sectional view (lower row), are as indicated in Fig. 1A, 1A’. The top view is a maximum intensity projection. Scale bar, 100 μm.

**Movie S16. Changes in surface geometry during the 1st cleavage division of a zebrafish embryo cultured in Ca^2+^-free E3 medium, related to Fig. 6F**.

Fluorescence movie of a Tg(*actb2:Utrophin-mCherry*) embryo cultured in Ca^2+^-free E3 medium showing labelled F-actin during the 1st cleavage division (41 - 63 mpf, 82 sec/frame). The different orientations, top view (upper panel) and xz-sectional view (lower panel), are as indicated in Fig. 1A, 1A’. The top view is a maximum intensity projection. Scale bar, 100 μm.

**Movie S17. Changes in surface geometry during the 1st cleavage division of a zebrafish embryo overexpressing CARhoA, related to Fig. 6G**.

Fluorescence movie of a Tg(*actb2:Utrophin-mCherry*) embryo injected with 600 pg CARhoA mRNA showing labelled F-actin during the 1st cleavage division (41 - 67 mpf, 56 sec/frame). The different orientations, top view (upper panel) and xz-sectional view (lower panel), are as indicated in Fig. 1A, 1A’. The top view is a maximum intensity projection. Scale bar, 100 μm.

