## Supplementary material for "Non-canonical cytokinesis driven by mechanical uncoupling via nematic flows and adhesion-based invagination": SI Theory Note

In this Supplemental Theory Note, we discuss two aspects of our modelling strategy: the first part discusses the active nematic gel theory for the actin cortex dynamics at the mesoscopic level, and the second part presents the theory of constriction at the macroscopic level.

### T1. ACTIVE GEL THEORY FOR ACTIN CORTEX DYNAMICS

We provide a comprehensive mathematical formulation and analysis of the active gel theory presented in the main text. We systematically build up complexity in the theoretical framework, beginning with the simplest isotropic model and progressively incorporating additional complexity to capture the essential features of experimentally observed actin dynamics. We also show how the additional assumptions we make are not only important for understanding detailed quantitative features of the data, but also fundamental qualitative features of the actin concentration and velocity fields.

We first examine an isotropic active gel formulation, then extend it to include nematic order parameters that enable anisotropic stress generation and shear flows, and finally introduce a two-component actomyosin model that distinguishes between actively contractile actomyosin networks (in the cable region, under the control of RhoA, see Fig. 2D, S2B) and passive ( low contractility) punctate actin structures in the poles of the cortex.

#### A. Isotropic active gel theory

We begin our analysis with an isotropic active gel model that treats the actin cortex as a viscous gel capable of generating internal contractile forces. In this formulation, we can still describe nematic alignment of the network, which is characterized by the traceless tensor $Q$ , but this local nematic order is simply passively aligned by the velocity field, without feedbacking to active stress generation. Thus, in this formulation, all active gel coefficients, and in particular the contractile force, remains isotropic regardless of local actin nematic direction.

The model couples three primary fields: actin density $\rho(x,t)$, velocity field $\mathbf{v}(x,t)$, and nematic order tensor $Q$ through the following system of equations:

$$\begin{aligned} \partial_t \rho + \nabla \cdot (\mathbf{v}\rho) &= -\alpha(\rho - \rho_0) + D\nabla^2 \rho \\ \gamma v_i &= \partial_j T_{ij} \\ T_{ij} &= 2\eta \tilde{e}_{ij} + \eta_b \partial_k v_k \delta_{ij} + \chi_0 \delta_{ij} \frac{2\rho}{\rho + \rho_0} \\ D_t Q_{ij} &= \frac{1}{\beta_2} H_{ij} + \beta_1 \tilde{e}_{ij} \end{aligned} \tag{T1}$$

with the material derivative $D_t Q_{ij} = \partial_t Q_{ij} + v_k \partial_k Q_{ij} + w_{ik} Q_{kj} + Q_{ik} w_{kj}$, which includes the advection by the velocity field $\mathbf{v}$ and the rotation by the vorticity tensor $w_{ij} = \frac{1}{2}(\partial_i v_j - \partial_j v_i)$ [1]. The traceless rate of strain tensor $\tilde{e}_{ij}$ is defined as:

$$\tilde{e}_{ij} = \frac{1}{2}\left(\partial_i v_j + \partial_j v_i\right) - \frac{1}{d}\partial_k v_k \delta_{ij} \tag{T2}$$

where $d$ is the spatial dimension. Lastly, the molecular field $H_{ij}$ in this simplest case describes the relaxation of the nematic order tensor $Q_{ij}$ to a uniform isotropic state, which is given by:

$$H_{ij} = -Q_{ij} + \kappa \nabla^2 Q_{ij} \tag{T3}$$

This system incorporates several key physical processes. The density equation includes advection by the velocity field, linear relaxation to equilibrium density $\rho_0$ via turnover with the cytoplasm at rate $\alpha$, and diffusion with coefficient $D$ (note that $D$ will be assumed as very small in this system, and can thus be seen as a small non-local flux in addition to the velocity field $\mathbf{v}$, [2]). Force balance operates in the low Reynolds number regime with friction coefficient $\gamma$. The stress tensor encompasses viscous contributions (shear viscosity $\eta$, bulk viscosity $\eta_b$) and active stress proportional to actin density with contractility parameter $\chi_0 > 0$, reflecting the contractile nature of the system. The saturating factor $\frac{2\rho}{\rho+\rho_0}$ prevents singularities at high actin densities [3]. The nematic order tensor evolves through advection, relaxation to the isotropic state with time-scale $\beta_2$, and flow alignment with strength $\beta_1$.

We note that the model presented in this section is the same as the one in Reymann *et. al.* [4] in the context of holoblastic cell division, where they showed that inward flow generated by the actin in the contractile ring is sufficient to align the nematic order along the ring.

#### 1. *Implementation of the model in 2D*

From here onwards, we restrict our attention to $d = 2$, as the cortical thickness is small compared to lateral dimensions. As discussed in Jülicher *et al* [1], integrating over the cortex thickness yields the same effective compressible 2D model. We note that in two dimensions, $2\partial_j \tilde{e}_{ij} = \partial_j \partial_j v_i$ as the other two terms cancel out. Thus the force balance equation simplies to:

$$\gamma \mathbf{v}_i - \eta \nabla^2 \mathbf{v}_i - \eta_b \partial_i (\nabla \cdot \mathbf{v}) = \chi_0 \partial_i \frac{2\rho}{\rho + \rho_0} \tag{T4}$$

In two dimensions, the traceless symmetric nematic tensor is parameterized using two scalar fields:

$$Q = Q_{xx} = -Q_{yy} \tag{T5}$$

$$q = Q_{xy} = Q_{yx} \tag{T6}$$

The evolution equations for these components are

$$\begin{aligned} \partial_t Q + v_x \partial_x Q + v_y \partial_y Q + (\partial_x v_y - \partial_y v_x) q &= \frac{\beta_1}{2}(\partial_x v_x - \partial_y v_y) - \frac{1}{\beta_2} Q + \frac{\kappa}{\beta_2} \nabla^2 Q \\ \partial_t q + v_x \partial_x q + v_y \partial_y q - (\partial_x v_y - \partial_y v_x) Q &= \frac{\beta_1}{2}(\partial_x v_y + \partial_y v_x) - \frac{1}{\beta_2} q + \frac{\kappa}{\beta_2} \nabla^2 q \end{aligned} \tag{T7}$$

All numerical simulations are performed with finite element method using the ngsolve library [5] with a system size of $1 \times 1$, with Dirichlet boundary conditions for the velocity field $\mathbf{v}$ and Neumann boundary conditions for $\rho, Q$ to ensure no flux at the boundaries. The initial condition is set to be $\rho = 1, \mathbf{v} = 0, Q = 0, q = 0$ everywhere unless otherwise stated. Additionally, we set $\kappa = 10^{-4}, D = 10^{-4}, \eta = 1, \eta_b = 0, \alpha = 1, \beta_1 = 1, \beta_2 = 1$ unless otherwise stated. Both $\kappa$ and $D$ are small compared to the system size since there is little diffusion, and are thus only included for numerical stability without key effects on the underlying dynamics. $\alpha, \eta$ are defined to be 1 by rescaling time and space as explained in the next section and we assume that $\beta_2$, the relaxation time scale of the nematic order, is of the same order of magnitude as the actin turnover time scale $\alpha^{-1}$, as we expect them to be dependent on similar biochemical mechanisms. The impact of flow on nematic order $\beta_1$ is difficult to estimate directly from experiments. Thus we set it to be 1 for simplicity, although we do note that varying $\beta_1$ by factors of 10 either way typically does not vary the result qualitatively.

We note that this is consistent with the best-fit values from Reynmann *et. al.* [4]: if we assume an actin-turnover time of 25s as measured in Arslan *et. al.* [6], these values correspond to $\beta_2 \alpha = 5.4, \beta_1 = 0.7$ (both in units of actin turnover time), which are of the same order of magnitude as the ones we use here.

#### 2. *Non-dimensionalization*

To identify the relevant parameter combinations governing system behavior, we first non-dimensionalize our equations, using characteristic length and time scales. A natural length scale is $l_0 = \sqrt{\eta/\gamma}$, commonly referred to as the hydrodynamic length [1] – in the force balance equation without the bulk viscosity term (or in 1D), the velocity field responds to density gradients with a screened-Coulomb-type Green's function with screening length of $l_0$. This length scale $l_0$ can vary widely between different systems - ranging from tens to hundreds of microns [4, 7–9] - as it depends on whether or not there is frictional contacts with a substrate. The natural time scale corresponds to the actin turnover time $t_0 = 1/\alpha$, which is typically tens of seconds in most systems [6].

We introduce dimensionless variables $x' = x/l_0$, $t' = t/t_0$, $v' = vt_0/l_0$, and $\rho' = \rho/\rho_0$ (when we later make $\rho_0$ vary in space, we rescale with the average value of $\rho_0$ in the poles of the cortex). We further note that we could also rescale $Q$ here to eliminate one of $\beta_1, \beta_2$, however, as we will see later, we will add a spontaneous phase separating term in $H_{ij}$ which will set the scale of $Q_{ij}$. Substituting into Eq. (T1) and relabeling the primed variables back to the original, we obtain the following.

$$\begin{aligned} \partial_t \rho + \nabla \cdot (\mathbf{v}\rho) &= -(\rho - 1) + \frac{D\gamma}{\eta\alpha}\nabla^2\rho \\ \mathbf{v}_i - \nabla^2 \mathbf{v}_i - \frac{\eta_b}{\eta}\partial_i(\nabla\cdot\mathbf{v}) &= \frac{\chi_0}{\eta\alpha}\partial_i\frac{2\rho}{\rho+1} \\ D_t Q_{ij} &= \frac{1}{\alpha\beta_2}(-Q_{ij} + \frac{\gamma\kappa}{\eta}\nabla^2 Q_{ij}) + \beta_1\tilde{e}_{ij} \end{aligned} \tag{T8}$$

This reveals that the isotropic system is characterized by six dimensionless parameters:

$$\frac{D\gamma}{\eta\alpha}, \quad \frac{\eta_b}{\eta}, \quad \frac{\gamma\kappa}{\eta}, \quad \frac{\chi_0}{\eta\alpha}, \quad \frac{1}{\alpha\beta_2}, \quad \beta_1 \tag{T9}$$

with the first three being small or unimportant in terms of resulting dynamics (as discussed above, actin diffusion is negligible in the experimental system as evidenced by the formation of depletion zones, and bulk viscosity effects are not prominent in our system). Furthermore, in terms of stress-exerting component, in the isotropic theory, $Q$ only reacts passively to the velocity field so we focus our current analysis on the dimensionless active stress parameter $\chi_0/\eta\alpha$.

#### *3. Model predictions and limitations*

With this isotropic model, we screen for parameters that can reproduce a depletion zone. We implement the contractile cable as a zone of higher carrying capacity $\rho_0(\mathbf{x})$ for actin, modeled as a top-hat function $H(\mathbf{x})$ centered at the origin with a width of 0.1 along the x direction and 0.8 along the y direction. Even with this very local activation of contractility, we find that we need to be in a very low hydrodynamic length regime, as convergent flows would propagate otherwise. This isotropic model successfully reproduces some key experimental features: increased actin density within the contractile cable, flanking depletion zones (since our theory is only at the linear level and non-linear effects will kick in when the density is close to zero, we satisfy ourselves with identifying regions of low density as depletion zones), outward flows in the transverse direction, and alignment of the nematic order in the actin cable, as shown in figure T2.

The mechanism underlying the depletion zone is as follows: the contractile force exerted by the actin cable contracts the nearby actin, creating a depletion zone on either side. Because the hydrodynamic length is small, the contraction does not propagate across the system, thus the bulk region maintains its original density and pulls actin on the other side of the depletion zone, creating an outward flow. Furthermore, due to the alignment of the nematic order with the shear rate of the flow, $Q$ is aligned with the y direction, as the flow on either side of the cable squeezes the actin perpendicular to the cable.

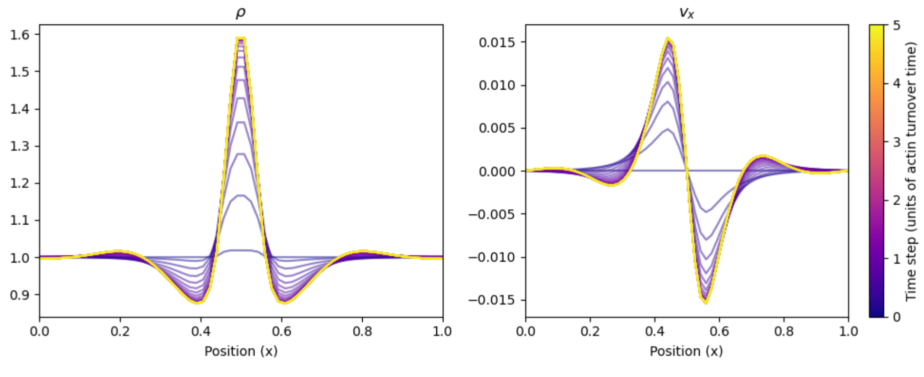


FIG. T1: Small hydrodynamic length limit $\sqrt{\eta/\gamma} = 0.1$ with system size of 1. $\chi_0 = 2.5$, $D = 5 \times 10^{-4}$ and $\rho_0(\mathbf{x}) = 1 + 0.2H(\mathbf{x})$ where $H(\mathbf{x})$ is a tophead function centered at the origin with a width of 0.1 along the x direction and 0.8 along the y direction.

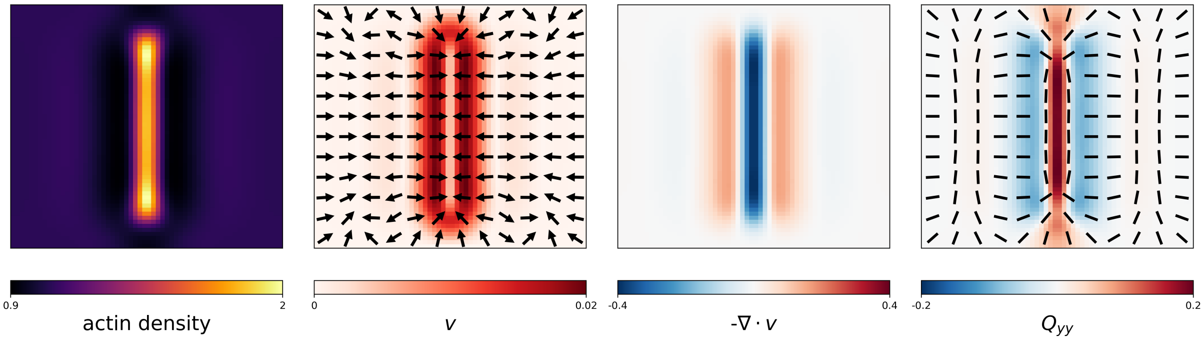


FIG. T2: 2D heatmap of $\rho, v, Q$ for the same parameters as in figure T1 at the final timepoint. The nematic order is aligned with the y direction inside the cable region due to the squeezing of the shear flow. We see inward flow close to the cable and outward flow outside the depletion zones.

However, the characteristics of the flow pattern contradicts the experimental observations (Fig. 2F in main text). The model generates purely outward transverse flows, whereas experiments demonstrate inward flow at cable termini combined with outward flow along cable flanks. This discrepancy arises from the fundamental constraint that isotropic active stress can only produce compressive or extensile flow configurations, precluding the generation of shear flows or anisotropic force distributions required to match experimental flow patterns.

A second limitation concerns the hydrodynamic length scale. The isotropic model requires small hydrodynamic length to generate the observed flow patterns. However, this implies large friction between cortex and surroundings, which would not have a clear mechanistic explanation, as supposed by measurements in [7–9]. In line with this, laser ablation experiments indicate that the hydrodynamic length is large and comparable to the system size (Fig. S4A, S4A', Movie S11). These findings demonstrate that isotropic active gel theory is fundamentally insufficient to capture the complete dynamics of the experimental system, motivating the extension to active nematic formulations - where the nematic order actually feedbacks on force generation.

### B. Nematic active gel theory

To incorporate anisotropic stress generation, we modify the active stress component of the force balance equation [1]:

$$T_{ij} = 2\eta\tilde{e}_{ij} + \eta_b \partial_k v_k \delta_{ij} + \chi_0 \delta_{ij} \frac{2\rho}{\rho + \rho_0} + \chi_1 Q_{ij} \frac{2\rho}{\rho + \rho_0} \tag{T10}$$

This formulation differs slightly from conventional active gel theories [1] by making the nematic stress term proportional to local actin density, ensuring that contractile forces arise only in regions of significant actin concentration. The parameter $\chi_1$ determines the strength and characteristics of nematic contractility: positive values correspond to contractile behavior (inward forces at aligned fiber ends), while negative values represent extensile behavior (outward forces at fiber ends). Based on measurements of recoil velocities in laser ablation experiments shown in Fig 4B, 4A, 4A', we expect the system to be strongly nematically contractile with $\chi_1 \gg \chi_0$.

The introduction of nematic stress creates a complex feedback between flow generation and nematic alignment. Initially, isotropic contraction generates inward flow that aligns the nematic order along the cable direction, consistent with previous isotropic predictions. The aligned nematic order then produces anisotropic stress that should, in principle, generate the experimentally observed flow pattern: inward-transverse flow on either side of the cable and outward flow at the termini as in experiments. However, we find that this mechanism faces a fundamental stability challenge. The shear flows generated by nematic stress act to destabilize the nematic order through the flow-alignment term ($\beta_1$ term) in Eq. (T1), creating a negative feedback that limits the magnitude of achievable nematic order and, consequently, the resulting shear flows.

This destablization can be demonstrated analytically in the low-friction limit, which appropriately describes our system, given that the hydrodynamic length is comparable to the system size, as discussed above. In this regime, assuming negligible advection and diffusion compared to other terms, at the linear level with $\rho = 1 + \delta\rho$ and $v, Q$

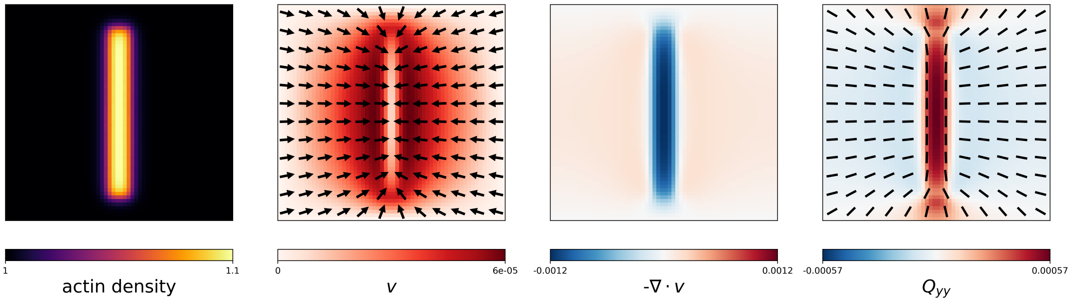


FIG. T3: Nematic active gel theory without any additional stablisation for the nematic order in the band. The parameters are $\chi_0 = 0.1, \chi_1 = 10, \rho(\mathbf{x}) = 1 + 0.1H(\mathbf{x})$.

small, the force balance equation and the nematic order equation in 1D simplify to:

$$\partial_x^2 v + \tilde{\chi}_0 \partial_x \delta\rho + \tilde{\chi}_1 \partial_x Q = 0 \tag{T11}$$

$$\frac{1}{2}\beta_1 \partial_x v - \frac{1}{\beta_2} Q = 0 \tag{T12}$$

where $\tilde{\chi}_0, \tilde{\chi}_1$ are the appropriately rescaled contractility paramaters $\chi_0, \chi_1$. Integrating the first equation with respect to $x$ yields

$$\partial_x v = -\tilde{\chi}_0 \delta\rho - \tilde{\chi}_1 Q + C \tag{T13}$$

where C is a constant of integration that ensures the Dirichlet boundary conditions for $v$ is satisfied. Substituting this into the second equation gives:

$$2\tilde{\chi}_1 Q = -\frac{\tilde{\chi}_0}{1 + \frac{1}{\beta_1 \beta_2 \tilde{\chi}_1}} \delta\rho + C \tag{T14}$$

where we have redefined C to absorb all factors in front of the constant term. Thus we have obtained an expression of $Q$ in terms of $\delta\rho$, allowing us to substitute this back into the first equation,

$$\partial_x v = -\delta\rho \frac{2\tilde{\chi}_0}{1 + \beta_1 \beta_2 \tilde{\chi}_1} + C \tag{T15}$$

Hence the effective contractility of the system must have the same sign as $\tilde{\chi}_0$. This implies that nematic order can only reduce the effective isotropic contractility of the system but cannot generate flows that are qualitatively different, as shown in figure T3. In other words, we cannot have outward transverse flow observed in experiments with only contractile nematics.

#### 1. *Stablisation of nematic order in actin cable*

Given the fundamental issue uncovered above of stabilization of the nematic order in the presence of nematic contractility, we explore the simplest extension of the model, which is a self-stabilizing mechanism that ensures that a non-zero nematic order is a locally stable configuration. In the equations, this is incorporated by modifying the molecular field $H_{ij}$ in Eq. (T3) to include a term that stabilizes the nematic order toward a position-dependent target magnitude $\bar{Q}(x)$, similar to the implementation of an activity induced isotropic-nematic transition in Cates *et. al.* [10]. We set $\bar{Q}(\mathbf{x}) = -1 + 2H(\mathbf{x})$ such that it is 1 inside the actin cable and -1 outside, representing that the actins inside the cable are stable once alignment is achieved, as evidenced by swift recovery of the nematic order after laser ablation (Fig. S3B, S3B', Movie S9) . The modified molecular field becomes:

$$H_{ij} = -(|Q|^2 - \bar{Q}(\mathbf{x})^2) Q_{ij} + \kappa \nabla^2 Q_{ij} \tag{T16}$$

We emphasize that we remain agnostic to the origin of this self-stablization mechanism, which could arise from biochemical feedback loops or other physical processes. One example is the so-called catch bond mechanism, where

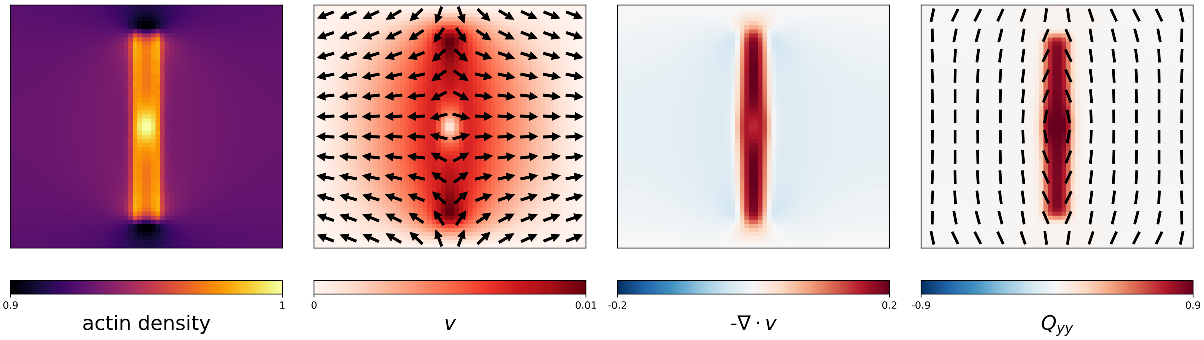


FIG. T4: Nematic active gel theory with self-stablization of nematic order in the coontractile region. The parameters are $\chi_0 = 0.1, \chi_1 = 0.3, \rho_0(\mathbf{x}) = 1 + 0.5H(\mathbf{x})$.

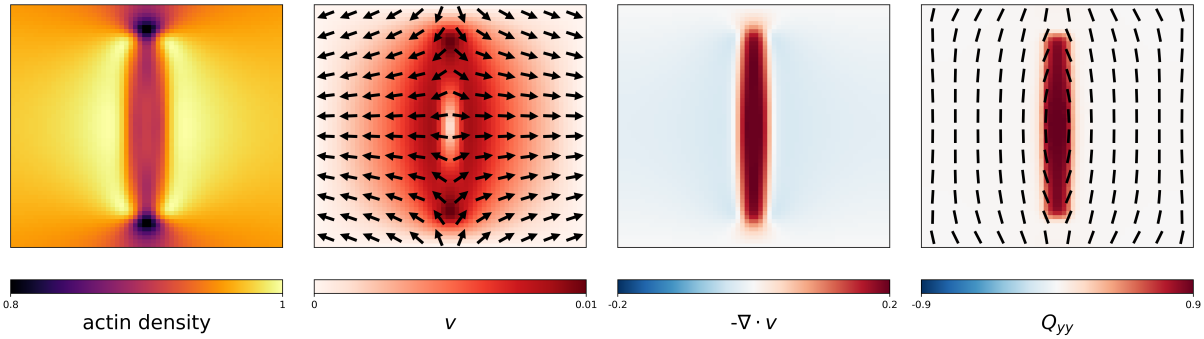


FIG. T5: Nematic active gel theory with self-stablization of nematic order in the coontractile region. The parameters are $\chi_0 = 0.1, \chi_1 = 0.3, \rho_0(\mathbf{x}) = 1 + 0.1H(\mathbf{x})$.

actomyosin contractility increases in response to applied mechanical stress, creating a positive feedback loop that stabilizes actin network under tension [11]. For instance, Ioratim-Uba et al. [12] demonstrated how such mechanochemical feedback can generate convergence extension flows in epithelial tissues similar to the ones observed in our system. When we implemented such mechanisms explicitly in our model, we found the outcome to be qualitatively similar to the one presented here, so in the spirit of minimum models, we choose to keep the current abstraction of the self-stabilization mechanism.

#### *2. Model Performance and Limitations*

In the biologically relevant regime in parameter space ($\chi_0 < \chi_1$), similar to the isotropic model, the initial isotropic contraction generates inward flow that aligns the nematic order along the cable direction (negative Q). This breaks the rotational symmetry of the system while simultaneously kicking the nematic dynamics away from the unstable fixed point at $Q = 0$. The aligned nematic order then produces anisotropic stress that generates the experimentally observed flow pattern: inward lateral flow on either side of the cable and outward flow at the ends as shown in panel 2 of figure T4. In contrast to before, the $\beta_1$ term is now dominated by the stablising effect and a non-zero steady-state nematic order is achieved, as shown in figure T4.

However, after a comprehensive parameter scan in $\chi_0, \chi_1, \beta_1, \beta_2$, the model fail to generate the observed depletion zone phenomena. If the carrying capacity in the contractile band $\rho_0(0)$ is set to be high, the system exhibits a region of enhanced density at the cable but no flanking depletion zones, as shown in figure T4. If the carrying capacity is set to be low, the actin cable gets advected away by the transverse flow and we have a region of reduced density (depletion zone) at the contractile region, as shown in figure T5. While it is possible, in extremely narrow region of parameter space, to achieve a stable actin cable with a slight drop-off of density on either side, the phenomena is not robust against small changes in parameters and thus we conclude that the nematic active gel theory with self-stabilization of nematic order is insufficient to capture the depletion zone phenomena observed in experiments.

#### C. Two-Component active gel theory

Previously, we have only treated the actin cortex as a single-component active gel, where the actin density $\rho$ represents the total concentration of all actin filaments. This brings limitations to the model: we assume that the actin turnover rate is uniform across the cortex and thus we cannot have the cable region with high actin density and the flanking regions in depletion simultaneously under the same velocity field. The assumption is quite unrealistic, as one could generically expect non-linearity, either in depolymerization or in polymerization, e.g. if the depletion zones are also depleted in actin assembly regulators, as shown in [2, 13].

To address these limitations, we extend the framework to incorporate distinct active and passive actomyosin populations. This two-component system captures the functional diversity of actin networks: in the contractile zone, actin co-localize with myosin to form contractile actomyosin, which generates active stress; while in the passive zone, punctate actin provides structural support without contributing to contractility.

##### 1. *Chemical Kinetics*

Let $\phi$ denote contractile actin density and $\psi$ passive actin density, which both are assembled from a common monomer pool with density $m$. The full chemical reaction network is described by:

$$\partial_t \phi = -k_1 \phi + k_2(\mathbf{x}) m \tag{T17}$$
$$\partial_t \psi = -k_3 \psi + k_4(\mathbf{x}) m \tag{T18}$$
$$\partial_t m = k_1 \phi + k_3 \psi - [k_2(\mathbf{x}) + k_4(\mathbf{x})]\, m - k_5 m + c \tag{T19}$$

where $k_1, k_2, k_3, k_4, k_5$ are reaction rates and $c$ represents constant monomer production. We model $k_2(\mathbf{x})$ and $k_4(\mathbf{x})$ as spatially varying functions that capture local assembly preferences – in the contractile zone, active actin assembly is favored ($k_4(\mathbf{x}) = 0$), while in the passive zone, punctate actin assembly is preferred ($k_2(\mathbf{x}) = 0$).

Assuming that monomer is always in large excess [14], the monomer density always stays around the steady state value of $m \approx c/k_5$. We tested that the model phenomena are unchanged qualitatively even if we relax this assumption. Under these conditions, the actin densities evolve according to:

$$\partial_t \phi = -k_1(\phi - \phi_0(\mathbf{x})) \tag{T20}$$
$$\partial_t \psi = -k_3(\psi - \psi_0(\mathbf{x})) \tag{T21}$$

where the spatial dependencies of $k_2, k_4$ have been absorbed into the equilibrium densities $\phi_0(\mathbf{x})$ and $\psi_0(\mathbf{x})$: $\phi_0(\mathbf{x}) = ck_2(\mathbf{x})/k_5 k_1$ and $\psi_0(\mathbf{x}) = ck_4(\mathbf{x})/k_5 k_3$. Phenomenologically, we simply write $\phi_0(\mathbf{x}) = H(\mathbf{x})$ and $\psi_0(\mathbf{x}) = 1 - H(\mathbf{x})$, where $H(\mathbf{x})$ is the top-hat function as defined before, to represent the distinct assembly preferences of the two actin populations inside and outside the contractile zone.

Adding back the advection, diffusion terms, together with the force balance and the nematic order equation, we obtain the two-component active gel model:

$$\begin{aligned}
\partial_t \phi + \nabla \cdot (\mathbf{v}\phi) &= -k_1 \left[\phi - \phi_0(\mathbf{x})\right] + D_\phi \nabla^2 \phi \\
\partial_t \psi + \nabla \cdot (\mathbf{v}\psi) &= -k_3 \left[\psi - \psi_0(\mathbf{x})\right] + D_\psi \nabla^2 \psi \\
\gamma v_i &= \partial_j T_{ij} \\
T_{ij} &= 2\eta \tilde{e}_{ij} + \eta_b \partial_k v_k \delta_{ij} + \chi_0 \delta_{ij} \frac{2\phi}{\phi + 1} + \chi_1 Q_{ij} \frac{2\phi}{\phi + 1} \\
D_t Q_{ij} &= \frac{1}{\beta_2} H_{ij} + \beta_1 \tilde{e}_{ij} \\
H_{ij} &= -(|Q|^2 - \bar{Q}(\mathbf{x})^2) Q_{ij} + \kappa \nabla^2 Q_{ij}
\end{aligned} \tag{T22}$$

where $D_\phi, D_\psi$ are the diffusion coefficients for the two actin populations, which we set to be very small as before. The force balance equation is similar to the nematic active gel theory, except that the active stress is now only generated by the contractile actin population $\phi$. Since $\phi$ is the active component, we rescale time with $k_1$ (same role as $\alpha$ in the single-component models) and hence set $k_1 = 1$ in simulations without loss of generality. Similar to before, we set the diffusion constants $D_\phi, D_\psi$ to be $10^{-4}$ as diffusion is small compared to advection.

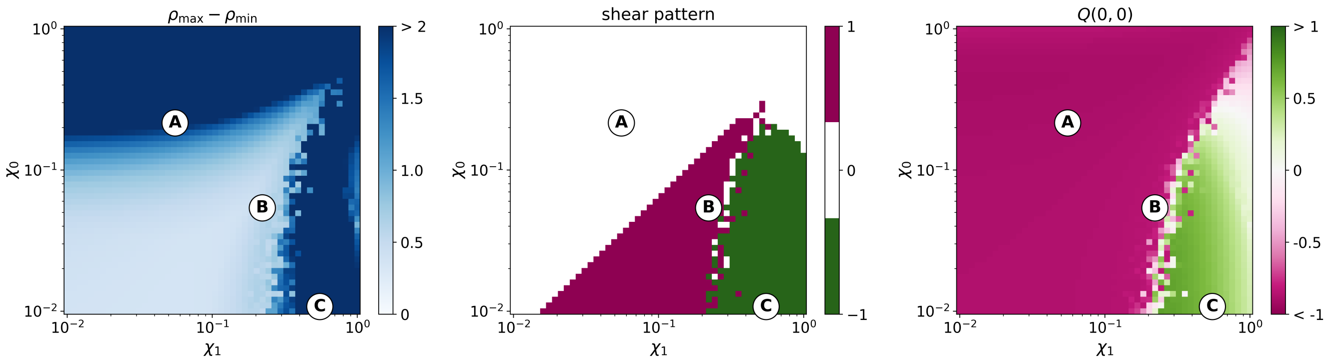


FIG. T6: Parameter sweep of the two-component active gel model in isotropic contractility $\chi_0$ and nematic contractility $\chi_1$. For all simulations, we set $\beta_1 = 1, k_3 = 0.2$. From left to right: the first panel shows the difference between the highest density and lowest density; the second panel shows the shear pattern with 0 identified with isotropic inward flow, 1 identified with shear flow with $\tilde{e}_{xx} > 0$ and -1 identified with shear flow with $\tilde{e}_{xx} < 0$; the third panel shows the nematic order $Q_{xx}$ at the centre of the actin cable, where positive values indicate alignment along the $x$ direction and negative values indicate alignment along the y direction. Examples of the three cases A-C are show in Fig. 3D.

A comprehensive parameter scan (figure T6) reveals that there is a large region in parameter space that qualitatively capture all key features of the experimental system: alignment of nematic order along the cable ($Q < 0$), inward flow at the cable ends, outward lateral flow on the sides (shear = 1), and depletion zones on either side of the cable (indicated by large $\rho_{\max} - \rho_{min}$ in the first panel). We note that the region with negative $Q$ arises because eventually the shear flow (inward at ends and outward at flanks) aligns the nematic order perpendicular to the cable direction, which is a much more stable configuration and we see indications of that in the pattern of myosin as shown in Fig. S3A.

A typical simulation in the operating parameter regime is shown in Fig. 3C and Fig. 3D: initially the enhanced contractility in the cable region generates inward flow that aligns the nematic order along the cable direction. The aligned nematic order then produces anisotropic stress that generates the experimentally observed flow pattern: inward flow along the cable and outward transverse flow. The contractile actin population $\phi$ is enhanced in the cable region due to the preferential assembly and high turnover rate, while the passive actin population $\psi$ is depleted in the same region for opposite reasons, creating a depletion zone on either side of the cable.

With an actin-turnover time of 25s as measured in Arslan *et al.* [6], the convergence phase in the upper panel of Figure 3E in the main text lasts for about 2 minutes, which is consistent with the experimental observations of 2 minutes in the lower panel of Figure 3E in the main text.

#### *2. CARhoA overexpression experiments*

Following discussions in the main text, overexpression of constitutively active RhoA (CARhoA) leads to reduced nematic order in the cable and consequently diminished outward transverse flows and actin depletion zones. Critically, the contractility $(\chi_1, \chi_0)$ remains unchanged, as discussed in the main text (Fig. 5B, 5C). Thus, we rationalize the CARhoA perturbation within the two-component active gel framework by reducing the value of $\bar{Q}^2$ in the cable. CARhoA overexpression is known to increase polymerization of actin filaments while preventing proper cycling of contractile functions [15]. There are a number of possible mechanisms for CARhoA overexpression to lead to an overall reduced nematic order: (1) enhanced growth of the actin filaments in confined environment leads to defects that disrupts the overall alignment order [16]; (2) increased nucleation of new filaments leads to a more branched network structure that's more difficult to align; (3) there are more cross-linking proteins binding to the filaments, which hinders their ability to align. Regardless of the exact mechanism, the net effect is an experimentally observed reduction in nematic order within the cable. On average, from experimental observations, the nematic order in the cable is reduced by about 39% upon CARhoA overexpression, corresponding to $\bar{Q}^2 \approx 0.6$. Once this Q parameter is fitted, given that we do not change any other parameters, we can still use the evolution of the flow fields and density profiles as tests of the model.

In simulations, we survey a range of $\bar{Q}^2$ values from 0.5 to 1 to mimic the spectrum of phenotypes as shown in Figure 5A of the main text. The results are shown in Figure 5E, 5E' with a typical simulation shown in Figure 5F.

### T2. THEORY OF CONSTRICTION OF THE ACTIN CABLE

The dynamics of division of a single cell into two daughter cells requires careful balance of forces between various components. Part 2 of the supplemental theory provides a detailed theoretical framework for understanding the mechanical aspects of this process, with particular emphasis on the forces generated by the actin cable and the cortex. Broadly, we coarsen the previous model by only considering effective surface and line tensions in the pole versus furrow regions, following the analytical arguments made in Turlier *et. al.* [17] and adapting them to meroblastic cleavages.

#### A. Phase 1 constriction

##### 1. Force balance

The cornerstone of our analysis is the force balance that governs the constriction process. We first write it with an effective energy description and minimize it to reach a stable force balance configuration. Three primary components contribute to the balance: the active energy from actomyosin contractility in the poles and cable $E$, the viscous dissipation that resists shape changes $D_{\text{vis}}$, and the energy injection from expanding microtubule overlapping zones $S$. These components are related through the energy balance equation:

$$\partial_t E + D_{\text{vis}} = S \tag{T23}$$

We will examine the three components of the equation in turn.

The active contractile energy $E$ arises from the deformation of the contractile cable and the actin cortex. Assuming that the system is well-modelled by two hemispheres - building on the physics of soap bubbles which have been shown to describe well cellular configurations [17], this energy is given by

$$E = L_f w(N_0^f - N_0^a) + 2A_p N_0^a \tag{T24}$$

where $L_f$ is the length of the furrow, $w$ is the width of the furrow, $N_0^f, N_0^a$ are tensions in the furrow and the cortex respectively, $A_p$ is the area of the pole. Let $R$ be the radius of the cortex and $\theta$ be the angle as indicated in the top panel of fig. T7, we further have $A_p = \pi R^2(1+\cos(\theta))$ due to volume conservation [17].

The rate of viscous dissipation $D_{\text{vis}}$ emerges from the viscous resistance of the actin cortex to deformation. By classical thermodynamics, viscous dissipation per unit volume is given by $\approx \eta(\partial_x v)^2$. Here, we use simple scaling arguments [17] to obtain,

$$D_{\text{vis}} = \frac{1}{2}\eta \left[ V_p \left( \frac{\dot{h_f}}{R} \right)^2 + V_f \left( \frac{\dot{h_f}}{h_f} \right)^2 \right] \tag{T25}$$

where $V_p = 2A_p e_p$ is the volume of the cortex at the poles and $V_f = L_f w e_f$ is the volume of the cortex at the furrow. $e_p$ and $e_f$ are the thickness of the cortex at the poles and the furrow respectively. $\eta$ is the actomyosin gel viscosity, as in the first part of the Theory Note.

Lastly, the injection of energy into the system due to expanding MT overlapping zones is given by

$$S = \alpha w(N_0^f - N_0^a) \tag{T26}$$

where $\alpha$ is a constant that parameterizes the expanding speed of the MT overlapping zones. Note that $\alpha t \neq L_f$ as the growth of the MT overlapping zone is not the only contributing factor to the length of the contractile cable – the contraction also plays a role in determining the length.

Having written down all the terms, we can see that we will eventually have a first order ODE with $(\theta, L_f)$ as the variables. What remains is geometrically relating $L_f$ to $\theta$. From the diagram in the top panel of fig. T7 and assuming the length of the MT overlapping zone grows linearly over time, we calculate the following:

$$\begin{aligned}
\phi &= \alpha t/(2R_0) \\
\Delta_1 &= R_0 \sin(\phi) \\
\Delta_2 &= h_f - R_0 \cos(\phi) \\
\tilde{R} &= \frac{\Delta_1^2 + \Delta_2^2}{2\Delta_2} \\
\psi &= \arcsin\Big(\Delta_1/\tilde{R}\Big) \\
L_f &= 2\psi\tilde{R}
\end{aligned} \tag{T27}$$

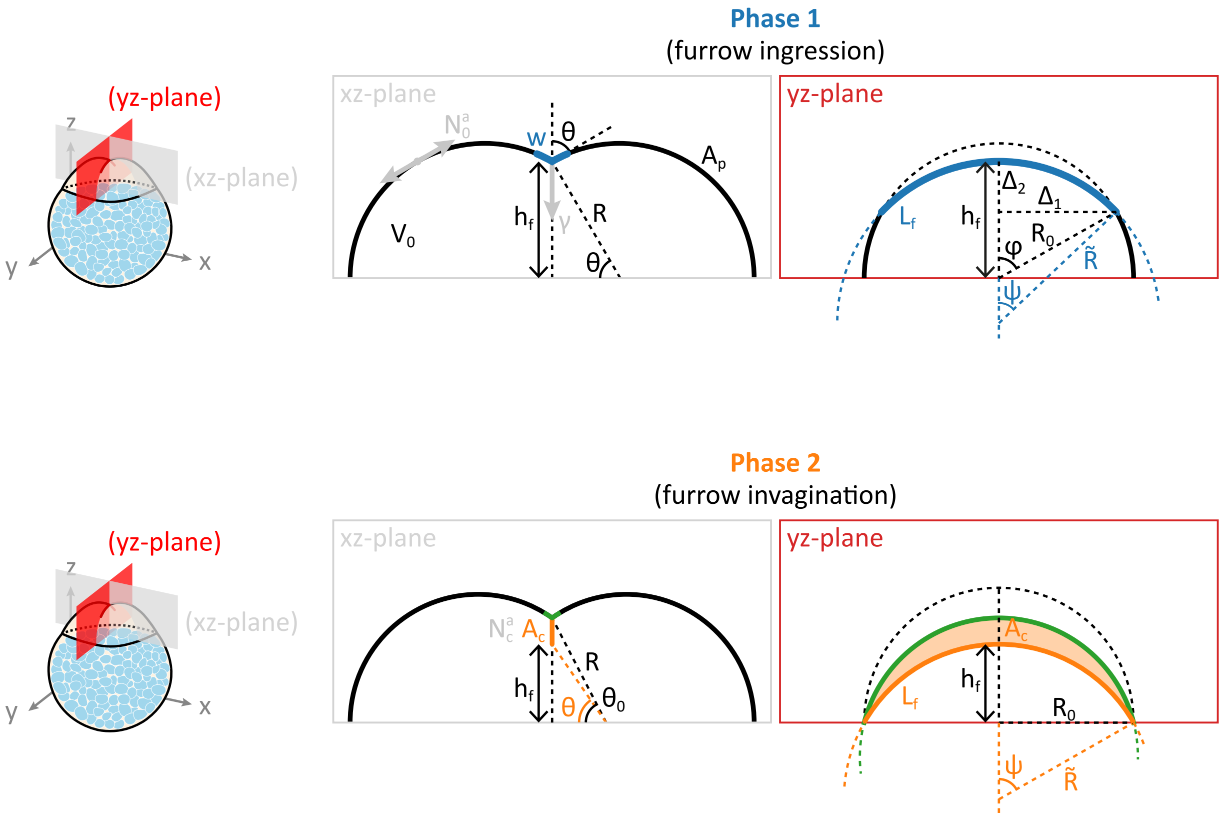


FIG. T7: Schematic illustration of the geometry of the constriction process in phase 1 and phase 2. In the top panel, the dark blue line represents the furrow. In the bottom panel, the orange line is the contractile cable, the green line is the and between the two line is the membrane-membrane contact area.

As the total volume of the embryo is conserved, we have an additional relation between $R$ and $R_0$: $R_0 = RF(\theta)^{1/3}$, where $F(\theta) = 1 + \frac{3}{2}\cos(\theta) - \frac{1}{2}\cos^3(\theta)$. Substiting this and $h_f = R\cos(\theta)$ into the previous set of equations, we obtain

$$\begin{aligned} \Delta_1/R &= F^{1/3}\sin(\phi) \\ \Delta_2/R &= \sin(\theta) - F^{1/3}\cos(\phi) \end{aligned} \tag{T28}$$

Now we have $L_f$ as a function of $\theta, \phi(t)$ as long as $R\sin(\theta) > R_0\cos(\phi) \to \sin(\theta)/F^{1/3} > \cos(\phi)$ .

#### 2. *Static energy comparison*

Having calculated $L_f$ as a function of $\theta, \phi(t)$, we can substitute it into the energy term to obtain $E$ as a function of $\theta, \phi(t)$ to see if contraction is energetically favorable.

Dividing the energy term $E$ by the total energy of the undivided cell $E_0 = 2\pi R_0^2 N_0^a$ , we obtain

$$\frac{E(\theta,\phi(t))}{E_0} = \frac{L_f(\theta,\phi(t))}{\pi R_0}\kappa + \frac{1+\cos(\theta)}{F(\theta)^{2/3}}, \qquad \kappa = \frac{w(N_0^f - N_0^a)}{2R_0 N_0^a} \tag{T29}$$

However, since the system can minimize its surface energy for free as MT overlapping zone expand, we also need to subtract this additional energy,

$$\frac{St}{E_0} = \frac{2\phi(t)w(N_0^f - N_0^a)}{2\pi R_0^2 N_0^a} = \frac{2\phi}{\pi}\kappa \tag{T30}$$

Putting them together, we obtain

$$\frac{E - St}{E_0} = \frac{\kappa}{\pi}\left(\frac{L_f}{R_0} - 2\phi\right) + \frac{1+\cos(\theta)}{F(\theta)^{2/3}} \tag{T31}$$

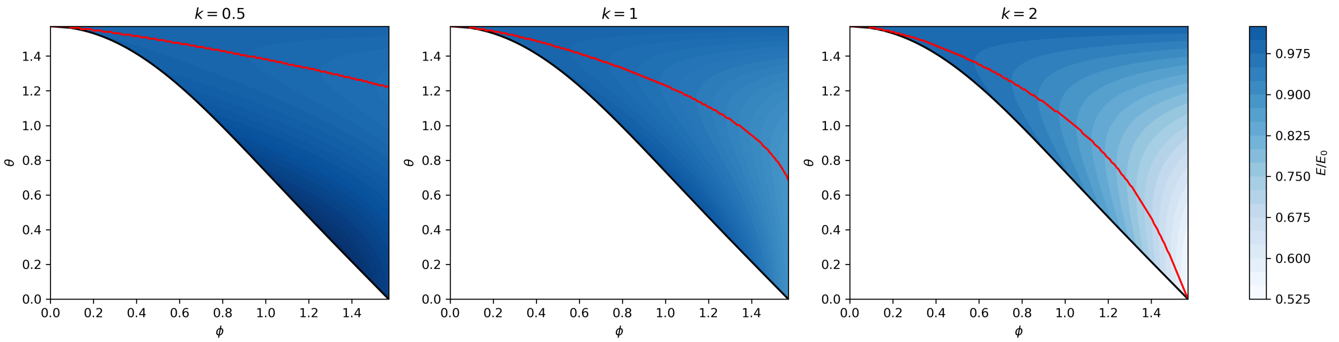


FIG. T8: Energy landscape of the system. Below the black line, the parameter regime is physically unfeasible as the furrow cannot be concave. The red lines indicate the energy minimum for each value of $\phi$.

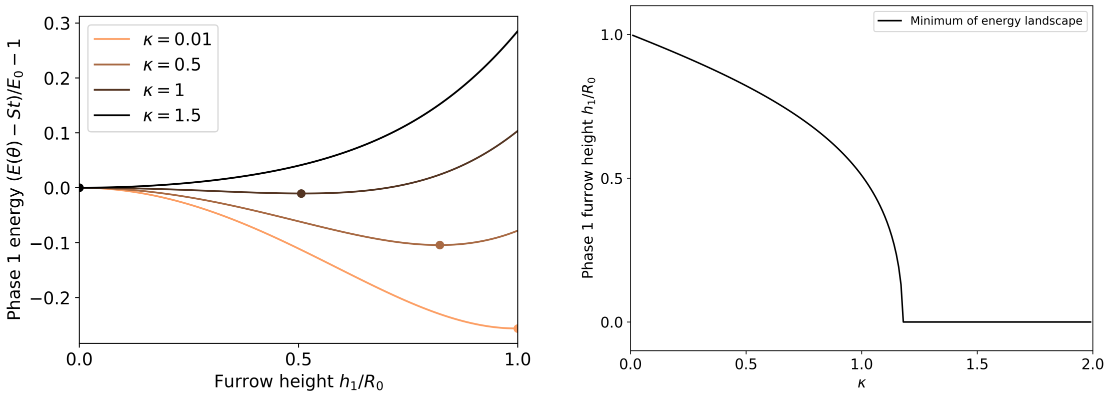


(a) Energy landscape as $\kappa$ varies

(b) Optimal height $h/R_0 = \sin(\theta)$ as $\kappa$ varies

FIG. T9: In the left panel, we show four examples of the energy landscape as $\kappa$ varies. The dots indicate the minima of the energies. In the right panel, we plot the optimal height as a function of $\kappa$, showing a continuous phase transition from the undivided state to the fully constricted state.

Note that the only parameter in this equation is $\kappa$, which corresponds to the ratio of the active energy in the furrow and the cortex. The energy landscape is shown in fig. T8. We can see that in physically feasible regime, as the furrow lengthens ($\phi$ increases) constriction follows ($\theta$ also decreases).

To determine the maximum possible extent of constriction in Phase 1, we only need to be concerned with when $\phi = \pi/2$, which corresponds to the MT overlapping zone reaching the margin of the blastodisc. At this point, the total energy is only a function of $\theta$ and the minimum of the energy yields the optimal constriction angle, as demonstrated in fig. T9. For small $\kappa$, the undivided state $\theta = \pi/2$ is more favorable. As $\kappa$ increases, the minimum of the energy smoothly shifts to smaller values of $\theta$, until the optimal angle reaches $\theta = 0$. As demonstrated in fig. T9b, in contrast to the first order phase transition observed in [17] in holoblastic cleavages, in our case, we have a continuous second order transition from the undivided state to the fully constricted state.

#### *3. Dynamics of constriction*

Next, we put back the dissipation term and look at the dynamics of the system. In the limit where constriction is much slower than the time to reach the steady state of the actin configuration, upper-bounded by the actin turnover rate in part 1, we can treat the embryo as a static system: for each value of $\phi$, the system relaxes to the $\theta$ that minimizes the energy (see fig. T8). In reality, the constriction speed is stalled by viscous friction. Starting with

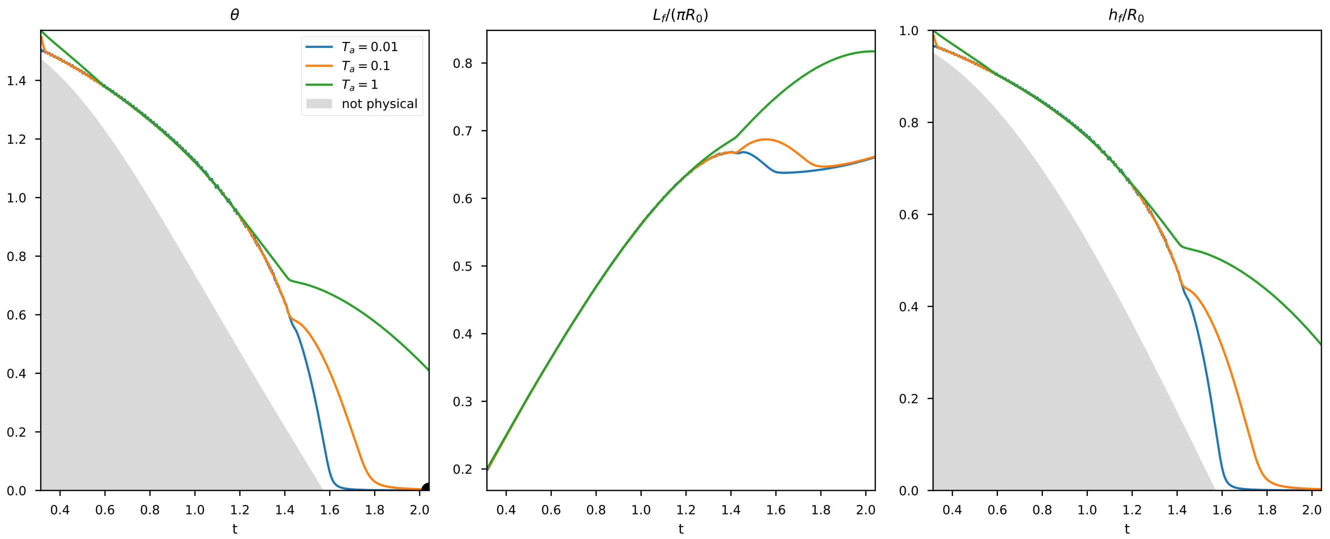


FIG. T10: Dynamics of constriction for various values of $T_a$. The black dot in panel 1 represents the minimum of the static energy. The rest of the parameters are set to $\kappa = 1.5, \lambda = 0.01, \dot{\phi} = 1$.

eq. T25, we also divide the term by $E_0$ and obtain

$$\begin{aligned}\frac{D_{\text{vis}}}{E_0} &= \frac{\eta e_p}{2N_0^a}\left[(1+\cos(\theta))\left(\frac{\dot{h}_f}{R_0}\right)^2 + \frac{we_f}{2R_0e_p}\frac{L_f}{\pi R_0}\left(\frac{\dot{h}_f}{h_f}\right)^2\right] \\ &= \frac{T_a}{2}\left[1+\cos(\theta)+\lambda\zeta\xi^{-2}\right]\dot{\xi}^2\end{aligned} \tag{T32}$$

where we have defined $\zeta(\theta,\phi) = L_f/(\pi R_0), \xi(\theta) = h_f/R_0 = \sin(\theta)/F^{1/3}, \lambda = \frac{we_f}{2R_0e_p}, T_a = \frac{\eta e_p}{N_0^a}$. Taking the time derivatives, we have

$$\frac{D_{\text{vis}}}{E_0} = \frac{T_a}{2}\left[1+\cos(\theta)+\lambda\zeta\xi^{-2}\right](\partial_\theta\xi)^2\dot{\theta}^2 \tag{T33}$$

Putting all the terms together in eq. T23 and denoting $\epsilon = E/E_0$, we obtain

$$\frac{T_a}{2}\left[1+\cos(\theta)+\lambda\zeta\xi^{-2}\right](\partial_\theta\xi)^2\dot{\theta}^2 + \partial_\theta\epsilon\dot{\theta} + \left(\partial_\phi\epsilon - \frac{2\kappa}{\pi}\right)\dot{\phi} = 0 \tag{T34}$$

This is a quadratic equation in $\dot{\theta}$ since we know $\dot{\phi}$ is just a constant. Overall, there are four parameters that control the constriction:

$$T_a = \frac{\eta e_p}{N_0^a}, \quad \dot{\phi} = \frac{\alpha}{2R_0}, \quad \lambda = \frac{we_f}{2R_0e_p}, \quad \kappa = \frac{w(N_0^f - N_0^a)}{2R_0N_0^a} \tag{T35}$$

The first two control the time scale of the constriction, the third is the ratio of volume of the furrow and the embryo, and the last one is the same $\kappa$ as before. Without loss of generality, we set $\dot{\phi} = 1$ by rescaling time. We further set $\lambda$ to be small since the width of the furrow $w$ is much smaller than the radius of the embryo $R_0$.

Generally, there are two solutions for $\dot{\theta}$ since equation (T35) is quadratic in $\dot{\theta}$. We choose the solution that has the same sign as $-\partial_\theta\epsilon$ to ensure that $\theta$ always go in the direction of decreasing energy, because the dissipation always goes in the direction that opposes the change. The solutions are shown in fig. T10 for $\kappa = 1.5$ corresponding to full constriction at the static level (fig. T9b). We can see that for as $T_a$ decreases from 1, meaning that the viscous timescale decreases relative to the expansion time of the MT overlapping zone, the fully constricted state is reached faster.

#### B. Phase 2 constriction

During phase 2, the actin cable detaches from the cortex and membrane adhesion protein is recruited at the membrane-membrane contact points as the cable constricts further. We hypothesize that the membranes that are stuck together has a lower surface tension compared to the rest of the cortex. Thus the total energy of the system is

$$E = L_f w(N_0^f - N_0^a) + 2A_p N_0^a + 2A_c N_c^a \tag{T36}$$

where the first term is the energy associated with the constricting cable as before, $A_c$ denotes the area of membrane-membrane contact as illustrated in the bottom panel of fig. T7 and $N_c^a$ is the surface tension of the membrane-membrane contacted area per membrane.

Since the geometry is slightly different in phase 2, we need to re-perform the computation for the length of the furrow $L_f$. The geometry is shown in fig. T7. Denoting $\theta_0$ as the angle at the end of phase 1 constriction, we have the following relations:

$$\begin{aligned} h_f &= \cos(\theta_0)\tan(\theta) \\ \tilde{R} &= \frac{R_0^2 + h_f^2}{2h_f} \\ \psi &= \arcsin\left(R_0/\tilde{R}\right) \\ L_f(\theta, \theta_0) &= 2\psi\tilde{R} \end{aligned} \tag{T37}$$

The contact area $A_c(\theta, \theta_0)$ can then be computed as the difference between the area under the arc at the end of phase 1 constriction and the area under the arc at the current position of the furrow, which we will not show explicitly here since it is a lengthy but straight-forward calculation. Putting together all the terms, the energy minus the source term is

$$\frac{E(\theta, \theta_0) - St_0}{E_0} = \frac{\kappa}{\pi}\left(\frac{L_f(\theta, \theta_0)}{R_0} - \pi\right) + \frac{1 + \cos(\theta_0)}{F(\theta_0)^{2/3}} + \frac{\nu}{\pi}\frac{A_c(\theta, \theta_0)}{R_0^2} \tag{T38}$$

where $\nu = \frac{N_c^a}{N_0^a}$. Now, in addition to $\kappa$, we have a new parameter $\nu$ that controls the energy landscape. As shown in [18, 19], membrane adhesion proteins can significantly lower the surface tension of the membrane-membrane contact zone, thus we expect $\nu < 1$.

We note that expression above is suitable for computing phase 1 energy in the limit where the contractile zone has extended to the margin ($\phi = \pi/2$), by simply setting $\theta_0 = \theta$. Three examples of the full energy landscape are shown in fig. T11 with $h_1$ as the upper point of the membrane-membrane contact and $h_2$ as the lower point. For the purpose of assessing the extent of constriction, only $h_2$ is relevant. The diagonal line corresponds to the total energy if only phase 1 constriction is considered. We can see that in cases where phase 1 mechanism is not sufficient to reach full constriction, the energy landscape is shifted downwards by the contribution of the membrane-membrane contact area. Interestingly, the overall minimum of the energy landscape typically has a higher $h_1$ than if only phase 1 constriction is considered, a result echoed by Fig. S6B. We refer to the main text for further comparisons with experiments. However, on the other hand, if phase 1 constriction is already sufficient to reach full constriction, as shown in the third panel of fig. T11, the addition of the membrane-membrane contact zone in fact hinders the constriction process by shifting the minimum to a higher value of $h_2$.

The full phase diagram in $\kappa{-}\nu$ space is shown in fig. T12. To compare with experimental data, we measured the final $h_1$ and $h_2$ values from 15 wild-type embryos and plotted the mean and standard deviation as contours in the two panels. As shown in fig. T12, there are a range of parameter values that could fit the $h_1$ and $h_2$ distribution independently. However, to fit both distributions simultaneously, we find that the best-fit parameters are $\kappa = 0.89, \nu = 0.72$, as indicated by the black cross in the left panel. Interestingly, the value of $\nu$ inferred here is consistent with previous measurements of the effect of E-cadherin on lowering surface tension [18, 19].

#### C. Perturbation experiments

Following discussions in the main text, we performed two perturbation experiments to test our theory on constriction – Calcium-free medium and CARhoA expression. In Calcium-free medium, the membrane adhesion is removed, corresponding to $\nu = 1$ (membrane-membrane contact is no longer energetically favorable). Since the tensile force in the furrow is unchanged, we keep $\kappa = 0.89$ as in wild-type. The results are shown in Fig. 6F, 6F'. For the CARhoA

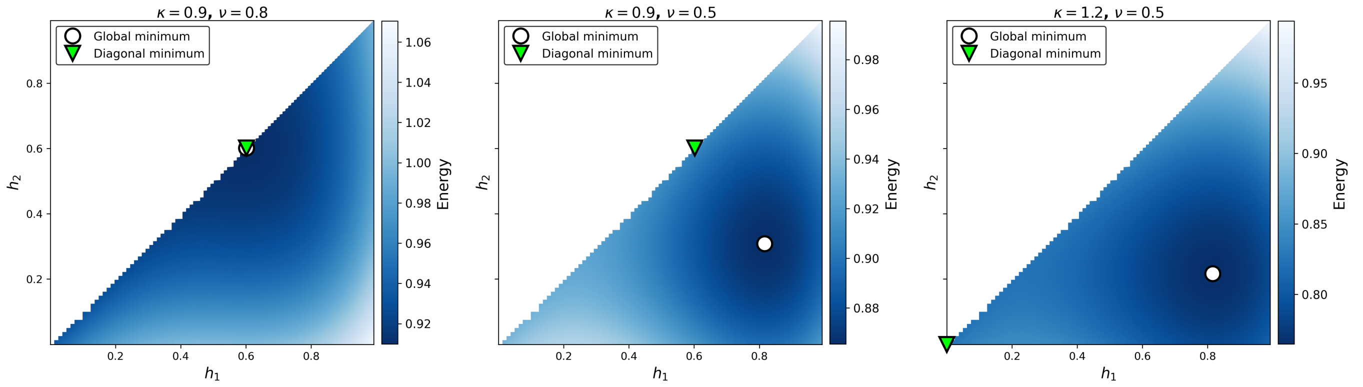


FIG. T11: Three examples of the energy landscape in phase 2. The diagonal corresponds to the energy landscape if only phase 1 constriction is considered. The green triangle indicates the minimum of the energy if only phase 1 is considered and the white circle represents the overall minimum.

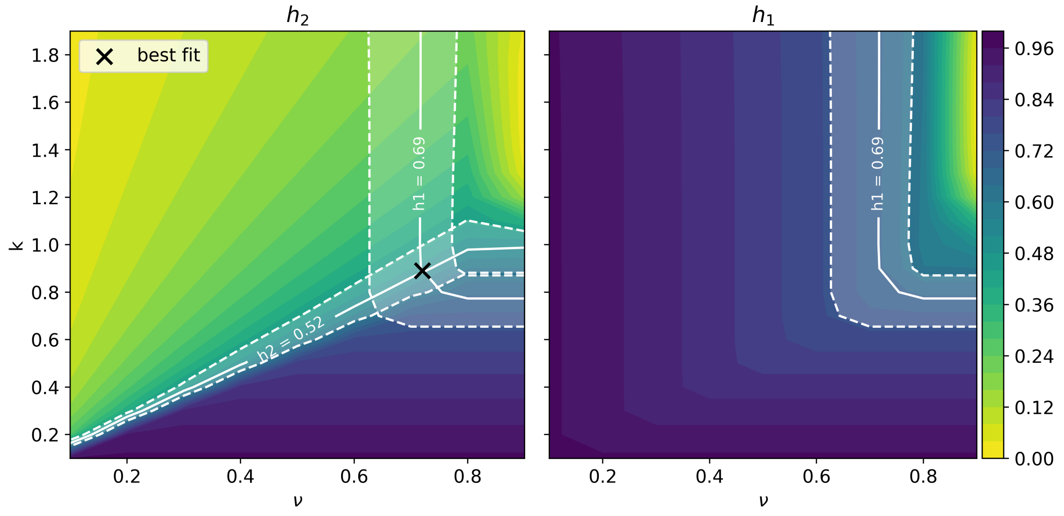


FIG. T12: Phase diagram of the constriction process including both phase 1 and phase 2 mechanisms in $\kappa - \nu$ space. Distribution of the final $h_1$ and $h_2$ values from WT constriction data are shown as contours in the two panels respectively. The best-fit parameters, indicated by the black cross in the left panel, corresponds to the intersection of the two contours.

overexpression experiment, the furrow does not detach from the cortex, inhibiting Phase 2 mechanisms. In the model, this is reflected by only limiting the constriction to Phase 1, and thus only $\kappa$ needs to be estimated. As discussed in section T1 C 2, CARhoA overexpression decreases the nematic order in the furrow, which in turn decreases the tensile force in the furrow by approximately 39%. Thus we set $\kappa = 0.55$. The results are shown in Fig. 6H, S6B.

---


[1] Frank Jülicher, Stephan W Grill, and Guillaume Salbreux. Hydrodynamic theory of active matter. *Reports on Progress in Physics*, 81(7):076601, 2018.

[2] Edouard Hannezo, Bo Dong, Pierre Recho, Jean-François Joanny, and Shigeo Hayashi. Cortical instability drives periodic supracellular actin pattern formation in epithelial tubes. *Proceedings of the National Academy of Sciences*, 112(28):8620–8625, 2015.

[3] Justin S Bois, Frank Jülicher, and Stephan W Grill. Pattern formation in active fluids. *Biophysical Journal*, 100(3):445a, 2011.

[4] Anne-Cecile Reymann, Fabio Staniscia, Anna Erzberger, Guillaume Salbreux, and Stephan W Grill. Cortical flow aligns actin filaments to form a furrow. *Elife*, 5:e17807, 2016.

[5] Joachim Schöberl. C++ 11 implementation of finite elements in ngsolve. *Institute for analysis and scientific computing, Vienna University of Technology*, 30, 2014.
[6] Feyza Nur Arslan, Édouard Hannezo, Jack Merrin, Martin Loose, and Carl-Philipp Heisenberg. Adhesion-induced cortical flows pattern e-cadherin-mediated cell contacts. *Current Biology*, 34(1):171–182, 2024.
[7] Mirjam Mayer, Martin Depken, Justin S Bois, Frank Jülicher, and Stephan W Grill. Anisotropies in cortical tension reveal the physical basis of polarizing cortical flows. *Nature*, 467(7315):617–621, 2010.
[8] Isabelle Bonnet, Philippe Marcq, Floris Bosveld, Luc Fetler, Yohanns Bellaïche, and François Graner. Mechanical state, material properties and continuous description of an epithelial tissue. *Journal of The Royal Society Interface*, 9(75):2614–2623, 2012.
[9] Daniel Boocock, Naoya Hino, Natalia Ruzickova, Tsuyoshi Hirashima, and Edouard Hannezo. Theory of mechanochemical patterning and optimal migration in cell monolayers. *Nature physics*, 17(2):267–274, 2021.
[10] ME Cates, SM Fielding, D Marenduzzo, Enzo Orlandini, and JM Yeomans. Shearing active gels close to the isotropic-nematic transition. *Physical review letters*, 101(6):068102, 2008.
[11] Wendy E Thomas, Viola Vogel, and Evgeni Sokurenko. Biophysics of catch bonds. *Annu. Rev. Biophys.*, 37(1):399–416, 2008.
[12] Aondoyima Ioratim-Uba, Tanniemola B Liverpool, and Silke Henkes. Mechanochemical active feedback generates convergence extension in epithelial tissue. *Physical review letters*, 131(23):238301, 2023.
[13] Aliaksandr A Halavatyi, Petr V Nazarov, Ziad Al Tanoury, Vladimir V Apanasovich, Mikalai Yatskou, and Evelyne Friederich. A mathematical model of actin filament turnover for fitting frap data. *European Biophysics Journal*, 39(4):669–677, 2010.
[14] William Brieher. Mechanisms of actin disassembly. *Molecular biology of the cell*, 24(15):2299–2302, 2013.
[15] Frank M Mason, Shicong Xie, Claudia G Vasquez, Michael Tworoger, and Adam C Martin. Rhoa gtpase inhibition organizes contraction during epithelial morphogenesis. *Journal of Cell Biology*, 214(5):603–617, 2016.
[16] Deb Sankar Banerjee, Simon L Freedman, Michael P Murrell, and Shiladitya Banerjee. Growth-induced collective bending and kinetic trapping of cytoskeletal filaments. *Cytoskeleton*, 81(8):409–419, 2024.
[17] Hervé Turlier, Basile Audoly, Jacques Prost, and Jean-François Joanny. Furrow constriction in animal cell cytokinesis. *Biophysical journal*, 106(1):114–123, 2014.
[18] Jean-Léon Maître, Hélène Berthoumieux, Simon Frederik Gabriel Krens, Guillaume Salbreux, Frank Jülicher, Ewa Paluch, and Carl-Philipp Heisenberg. Adhesion functions in cell sorting by mechanically coupling the cortices of adhering cells. *science*, 338(6104):253–256, 2012.
[19] Jean-Léon Maître, Ritsuya Niwayama, Hervé Turlier, François Nédélec, and Takashi Hiiragi. Pulsatile cell-autonomous contractility drives compaction in the mouse embryo. *Nature cell biology*, 17(7):849–855, 2015.
